# Transient reprogramming limits AP-1-associated chromatin opening and transposable element activation to preserve hematopoietic stem cell function during aging and stress

**DOI:** 10.64898/2026.08.28.747860

**Authors:** Audrey Porquet, Mathieu Bohm, Hassan Ait-Ougouram, Thu-Hà Trinh, Rabie Chelbi, Mengliang Ye, Ollivier Milhavet, Jean-Marc Lemaître, Nathalie Droin, Elina Zueva, Catherine M. Sawai, Emilie Elvira-Matelot, Françoise Porteu

**Affiliations:** University Paris-Saclay, Gustave Roussy, INSERM U1287, Hematopoietic stem cells and the development of myeloid malignancies, F-94805, Villejuif, France; INSERM Unit 1312 BRIC, University of Bordeaux, Bordeaux, France; Inovarion, 75005 Paris, France; INSERM U932, Institut Curie, Paris Sciences-et-Lettres Research University, 75005 Paris; IRBM, University of Montpellier, INSERM, CNRS UMR1183, Montpellier, France; University Paris Saclay, Gustave Roussy, INSERM, Fundamental genomics Platform, F-94805, Villejuif, France

## Abstract

Hematopoietic stem cell (HSC) aging is associated with epigenetic remodeling, yet the molecular mechanisms driving these changes, their overlap with stress-induced alterations, and whether this course can be durably reset remain incompletely understood. Here, we show that transient induction of the Yamanaka factors OCT4, SOX2, KLF4, and MYC in young mice durably delays and partially reverses physiological and LPS-driven HSC aging in mice. Transient reprogramming improved hematopoietic reconstitution, reduced myeloid bias, and limited DNA damage. Multi-omic analyses revealed reduced chromatin accessibility at AP-1-enriched regulatory regions, attenuated age-associated AP-1 transcriptional programs, and repression of transposable elements (TEs). Pharmacological AP-1 inhibition prevented LPS-induced TE activation and loss of HSC clonogenicity. Reverse transcriptase inhibition in aged mice reduced DNA damage and improved HSC function, demonstrating a functional contribution of TE activity to HSC decline. Together, these findings identify AP-1-associated chromatin remodeling as a candidate mechanism linking inflammatory stress, TE activation and HSC aging.

## Introduction

Hematopoietic stem cells (HSCs) sustain lifelong blood cell production. However, their functional properties are profoundly altered during aging. In particular, aged HSCs exhibit a marked decline in repopulation capacity and a differentiation bias toward the myeloid lineage, resulting in increased myeloid cell production and a progressive deterioration of adaptive immune responses ^1, 2^. These age-associated changes are accompanied by an expansion of the phenotypic HSC compartment and a remodeling of the HSC pool, characterized by a substantial increase in myeloid- and platelet-biased HSCs, which predominantly generate myeloid cells, and a comparatively smaller expansion of lymphoid-biased HSCs, which preferentially give rise to lymphoid lineages. Aged HSCs also accumulate DNA damage and acquire somatic mutations as a consequence of environmental stress and replication-associated errors. Although most of these mutations are functionally neutral, some of them confer a selective clonal advantage, leading to the expansion of mutant clones over non-mutated cells with age. This phenomenon, known as clonal hematopoiesis of indeterminate potential, occurs in otherwise healthy individuals and is associated with an increased risk of developing myeloid malignancies^3^.

HSC aging results from both cell-intrinsic alterations and extrinsic environmental changes, including chronic inflammation and dysfunction of the bone marrow niche that supports HSC maintenance^1, 2^. However, transplantation experiments and prolonged exposure of aged HSCs to a young bone marrow environment are insufficient to fully restore their functional defects^4^, underscoring the predominant role of intrinsic mechanisms in HSC aging. These observations suggest that strategies directly targeting the molecular and cellular features of aged HSCs may be necessary to delay or reverse HSC aging and preserve long-term hematopoietic function.

Numerous cell-intrinsic regulators of HSC aging have been identified, including impaired DNA repair leading to the accumulation of genomic damage, epigenetic, and transcriptional dysregulation, altered cell polarity, and defective mitochondrial activation and metabolism ^1, 2, 5–10^. Recent studies have particularly highlighted the central role of epigenetic alterations in aging^8, 10–12^. One hallmark of aging across multiple tissues is the widespread destabilization of heterochromatin, which results in the derepression and overexpression of transposable elements (TEs) ^13, 14^. TEs, which account for nearly half of the human and mouse genomes, include long terminal repeat (LTR) elements such as endogenous retroviruses (ERVs), as well as non-LTR elements including long and short interspersed nuclear elements (LINE-1/L1 and SINEs). Long considered “junk DNA,” TEs have recently emerged as important contributors to aging and cancer, particularly in contexts associated with chromatin disorganization. We and others have demonstrated that TE expression is increased in aged HSCs and in several models of premature aging induced by irradiation, chemotherapy, or chronic inflammation^13, 15–20^. We further showed that the reduction of the heterochromatin mark H3K9me3 observed in HSCs following irradiation or chronic inflammatory stress occurs predominantly at evolutionarily recent and active LINE-1 and LTR elements (L1Md and IAP in mice, respectively) ^19, 20^. This epigenetic erosion contributes to persistent DNA damage and long-term HSC dysfunction, both of which can be partially reversed by reverse transcriptase inhibitors (RTIs) ^15, 19, 20^. Together, these findings suggest that stress- or age-induced heterochromatin alterations and TE derepression may constitute a common molecular mechanism driving HSC dysfunction. However, the causal contribution of heterochromatin disorganization and TE activation to physiological HSC aging remains to be formally established. Moreover, although repeated inflammatory or replicative stress induce HSC defects resembling those observed during aging, the molecular links between stress responses and aging, and whether modulation of these pathways could rejuvenate HSC function remains unknown.

Recent studies have shown that senescent fibroblasts from centenarians, as well as aged HSCs, reprogrammed into induced pluripotent stem cells (iPSCs) using the Yamanaka factors OCT4, SOX2, KLF4, and MYC (OSKM), reacquire youthful characteristics upon redifferentiation into their somatic lineages^21, 22^. More recently, transient cellular rejuvenation without loss of somatic identity has been achieved in vitro through short-term expression of OSKM factors. In vivo, partial reprogramming induced by repeated cycles of transient OSKM expression have been shown to extend lifespan in premature aging mouse models and ameliorate aging phenotypes across multiple tissues^23–27^. These effects include restoration of tissue architecture, rejuvenation of DNA methylation and H3K9me3 patterns, resetting of epigenetic clocks, and improved regenerative capacity in both premature and physiological aging models. Collectively, these studies highlight the close relationship between rejuvenation and pluripotency programs and underscore the major role of epigenetic remodeling in aging. However, the molecular mechanisms underlying rejuvenation induced by transient reprogramming, particularly the contribution of TEs to this process, remain largely unknown. In addition, the impact of transient reprogramming on tissue stem cell maintenance during physiological aging, as well as its potential effects on HSC stress resilience has not yet been investigated.

Here, we show that a single short induction of OSKM factors early in life is sufficient to improve bone marrow and peripheral blood parameters, enhance HSC reconstitution capacity, restore the lymphoid-to-myeloid differentiation balance, and reverse many of the molecular alterations characteristic of aged HSCs. Early transient reprogramming prevents age-associated heterochromatin loss and chromatin opening at TEs and at binding sites for the stress-response transcription factor activator protein-1 (AP-1), thereby limiting TE expression, TE-induced DNA damage, and HSC functional decline. Transient OSKM induction also enhances the resilience of young HSCs to chronic inflammatory stress. These findings suggest that aging is associated with a persistent memory of stress-induced epigenetic alterations in HSCs and raise the possibility that limiting TE derepression and chronic AP-1 activation may represent a strategy to rejuvenate HSCs.

## Results

### Partial transient reprogramming early in life delays age-induced myeloid bias and HSC defects

To gain insight into the key intrinsic mechanisms underlying HSC aging, we hypothesized that these mechanisms would be among the cellular processes reversed during functional rejuvenation. To test this hypothesis, we first assessed whether physiological HSC aging could be slowed using an in vivo model of partial transient reprogramming. We used double heterozygous Rosa26^rtTA/+^; Col1a1^4F2A/+^ transgenic mice to induce controlled expression of the four Yamanaka transcription factors (4TF mice) through an rtTA transactivator upon doxycycline (Dox) administration in the drinking water^23, 26^ (Extended Data Fig. 1A). To assess the long-term persistence of the expected rejuvenating effects associated with partial reprogramming, we adopted a simplified protocol that we had previously described. ^23^ Rather than applying repeated treatments throughout life, reprogramming factors were induced only once using a low dose of Dox for 16 days at 2-3 months of age (Fig. 1A). Mice were then left untreated and analyzed at 20–24 months of age. Aged reprogrammed mice treated with Dox (O-Dox) were compared with age-matched untreated mice (O-NT) and with young (3-4 months) untreated mice (Y-NT).

**Figure 1:**
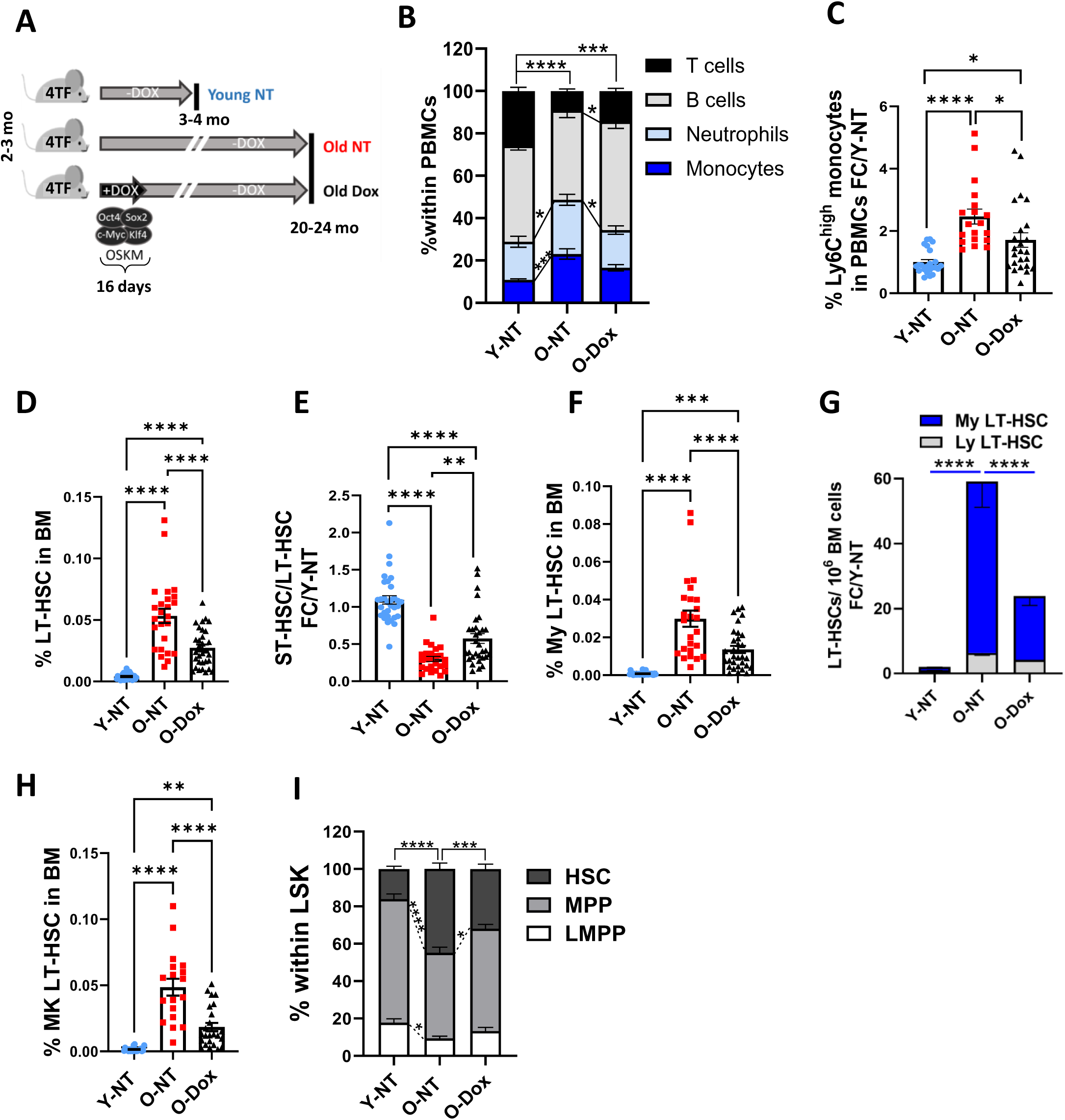
Partial reprogramming early in life prevents phenotypic changes associated to aging in blood and BM compartments. **(A)** Partial reprogramming induction protocol: young (2-3 months old) *R26 Col1a1* (4TF) mice are treated during 16 days with Dox (0.5mg/mL) diluted into the drinking water. Blood and BM aging hallmarks are analyzed at 20 -24 months of age in treated (O-Dox), non-treated (O-NT) and young (Y-NT) mice. (B-I) Flow cytometry analysis. **(B)** Repartition of myeloid and lymphoid cells; **(C)** Frequency of proinflammatory monocytes (CD11b Ly6G Ly6C), normalized to the Y-NT group; **(D)** Frequency of LT-HSCs in the BM; **(E)** Ratio of absolute numbers of ST-HSC/LT-HSC in the BM normalized to Y-NTgroup; **(F)** Frequency of myeloid-biased (My) LT-HSCs (CD150); **(G)** Numbers of myeloid- and lymphoid (Ly, CD150^low^)-biased LT-HSCs per million BM cells, normalized to the Y-NT group; **(H)** Frequency of megakaryocytic (MK)-biased LT-HSCs (CD41); **(I)** Repartition of HSCs, multipotent progenitor (MPP) and lymphoid multipotent progenitor (LMPP) within LSK cells. B, G, I: 19 to 31 mice per group. Two-way ANOVA with multiple comparisons test. Means +/- SEM. C-F, H: Each dot represents an individual mouse. One-way ANOVA with multiple comparisons test. Adjusted p-values are reported as: * (p < 0.05), **(p < 0.01), ***(p < 0.001), and ****(p < 0.0001).

We first assessed whether OSKM factors were expressed in HSCs following Dox treatment and whether this expression was maintained over time. RT-qPCR analysis showed that all transcription factors (TFs) are expressed or displayed increased expression in HSCs (defined as Lin^−^Sca1^+^Kit^+^ CD34^−^Flk2^−^ or LSK CD34^-^Flk2^-^) at the end of the treatment period (day 16) (Extended Data Fig. 1B). This expression was transient, as *Oct4* and *Sox2* were no longer detectable one month after treatment, while *Klf4* and *Myc* expression returned to levels comparable to those observed in untreated mice.

Analysis of blood parameters across the three groups of mice showed that Dox treatment early in life was sufficient to restore red blood cell and platelet counts to youthful levels (Extended Data Fig. 1C, D) and to reduce both the myeloid bias and the increase in pro-inflammatory monocytes associated with aging (Fig. 1B, C). In the bone marrow (BM), consistent with previous reports^1, 2^, the number of phenotypic long-term (LT)-HSCs (LSK Flk2^−^CD34^-^CD150^+^CD48^−^) markedly increased with age (Fig. 1D, Extended Data Fig. 1E), whereas the ratio of short-term (ST)-HSCs (LSK Flk2^−^CD34^+^CD150^+^CD48^−^) to LT-HSCs decreased (Fig. 1E), indicating the early loss of differentiation capacity that characterizes HSC aging. Early-life Dox treatment partially corrected these defects. Similarly, the age-associated increase in myeloid-biased HSCs (CD150^hi^ LT-HSCs) ^28, 29^, which account for most of the expansion of the HSC compartment in aged BM (Fig. 1F, G), as well as the increase in megakaryocyte-biased HSCs (CD41^+^ LT-HSCs, Fig. 1H), was delayed following Dox treatment. Finally, Dox treatment partially restored near to youthful levels the altered proportions of HSCs relative to myeloid- and lymphoid-primed multipotent progenitors observed during aging (Fig. 1I). Importantly, Dox administration to 2-month-old WT C57BL/6 mice had no effect on any of these parameters (Extended Data Fig.2 A-C), demonstrating that the observed effects were specific to OSKM induction and required transgene expression. Moreover, separate analyses of blood and BM in male and female 4TF transgenic mice revealed comparable rejuvenating effects following transient partial reprogramming (Extended Data Fig. 2D, E). Both sexes were therefore included in all subsequent experiments.

**Figure 2:**
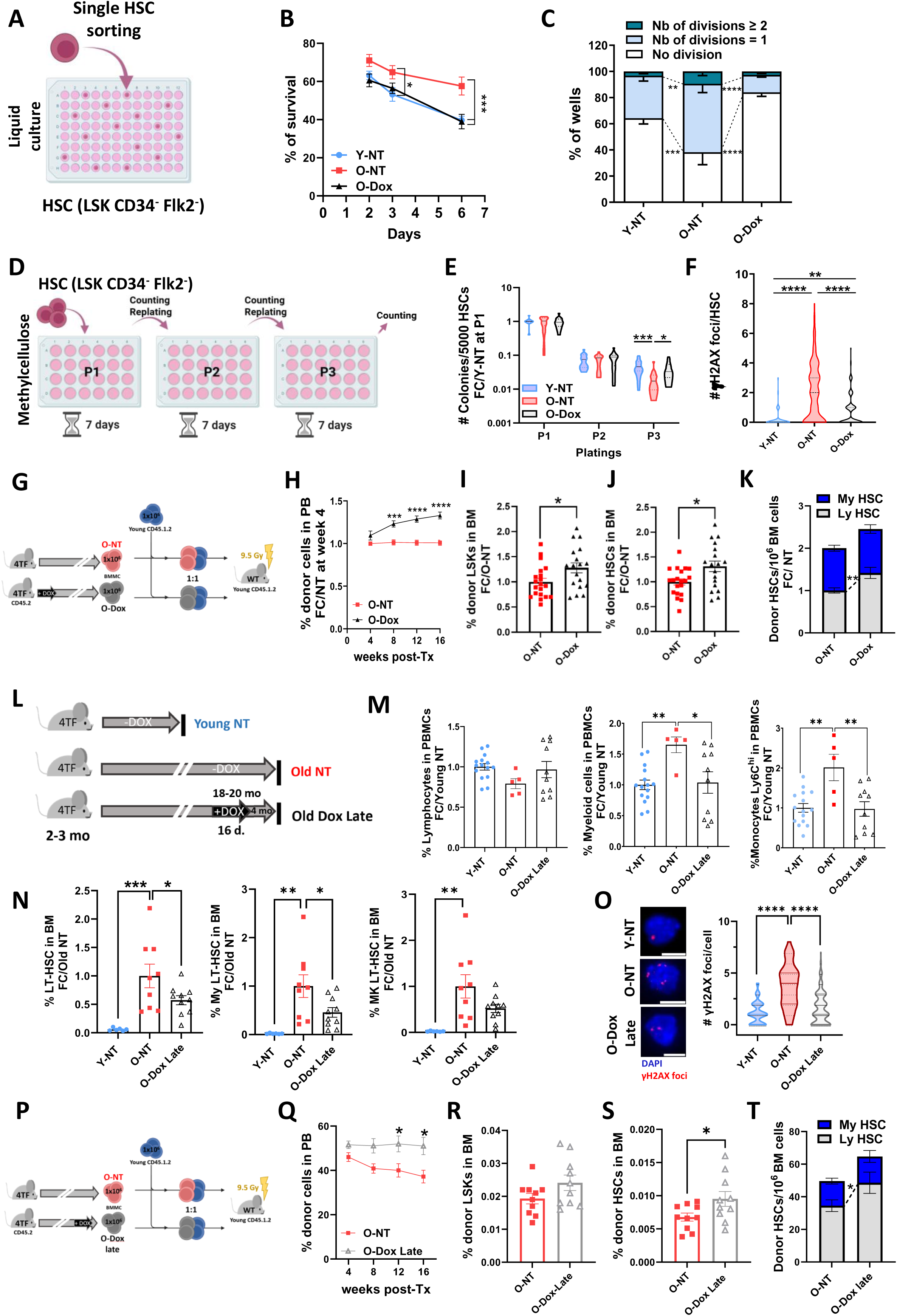
Early and late partial reprogramming prevents age-associated loss of HSC function. **(A-C)** Single HSC liquid cultures. (A) Experimental design; (B) Survival, defined as the percentage of wells containing at least one cell at the indicated time points. Y-NT (n=11), O-NT (n=8), O-Dox (n=10) from 3 independent experiments; (C) Repartition of wells containing 1 (quiescent), 2 (one division) or >2 (more than 1 division) HSCs after 48h of culture. Means +/-SEM. Y-NT (n=5), O-NT (n=4) and O-Dox (n=5) from 2 independent experiments. **(D, E)** CFU serial replating assay in methylcellulose. Experimental design (D) and numbers of colonies generated from 5000 HSCs (E). Y-NT (n=16), O-NT (n=12), O-Dox (n=14) mice from 5 independent experiments. Violin plots presenting median and quartiles. **(F)** γH2AX foci quantification by IF in HSCs. Y-NT (n=2) O-NT (n=4), O-Dox (n=4); 2 independent experiments. Violin plots presenting median and quartiles. **(G-K)** Competitive BM transplantations. (G) Experimental design; (H) Percentage of donor 4TF mice-derived cells (CD45.2+) in peripheral blood at different times after transplantation. Means +/- SEM, O-NT (21), O-Dox (18) from 4 independent experiments, normalized to NT group at week 4 post-transplantation; (I) Frequency of donor LSKs and (J) HSCs in the BM of recipients 16 weeks following transplantation, normalized to the NT group. Each dot represents a mouse. (K) Donor My and Ly-biased HSCs defined as LSK CD34^-^ Flk2^-^ CD150^low/high^ per one million of BM cells, normalized to the NT group. Means +/- SEM. **(L-T)** Partial reprogramming induced in aged (18-20 mo) 4TF mice (O-Dox late mice). (L) Experimental design; (M) Blood and (N) BM analysis in Y-NT, O-NT and O-Dox Late 4TF mice. Data are normalized to Y-NT (M) or O-NT groups (N). Means +/- SEM. One dot represents a mouse; (O) γH2AX foci representative images and quantification in HSCs. Y-NT (n=2), O-NT (n=1), O-Dox Late (n=3). 2 independent experiments. At least 50 HSCs per condition per experiment. Violin plots presenting median and quartiles; (P-T) Competitive transplantations with BM from O-NT and O-Dox late mice. (P) Experimental design; (Q) Donor chimerism in the blood over time. Means +/- SEM. O-NT (n=10), O-Dox (n=10) from 2 independent experiments. (R) Frequency of donor LSKs and (S) HSCs in the BM of recipients at 16 weeks. Each dot represents a mouse. (T) Donor My- and Ly biased HSCs per million of BM cells. Means +/- SEM. B, C, E, H, K, Q, T: Two-way ANOVA with multiple comparisons test. F, M, N, O: One-way ANOVA with multiple comparisons test. Adjusted p-values are reported as: * (p < 0.05), **(p < 0.01), ***(p < 0.001), and ****(p < 0.0001). I, J, R, S: T-test. * pval<0.05.

Single-cell in vitro liquid cultures of HSCs showed that the age-associated increase in HSC survival and loss of quiescence, as measured by the number of wells containing 0, 1 or more than 1 cell, respectively, after 48 h of culture—was completely abolished following early-life Dox treatment (Fig. 2A-C). Partial reprogramming also restored HSC self-renewal capacity, as assessed by sequential clonogenic replating assays (Fig. 2D, E). Untreated aged HSCs showed increased γH2AX foci as compared to both young and age-matched Old Dox-treated HSCs (Fig. 2F and Extended Data Fig. 3A), suggesting that Dox prevented the age-associated increase in DNA damage which is linked to loss of HSC function.

To further evaluate HSC function, we first performed non-competitive serial transplantation experiments using BM from young, 20-month-old O-NT, and O-Dox mice (Extended Data Fig. 3B). In primary and secondary transplantations, all recipient mice were successfully reconstituted; however, donor chimerism in peripheral blood was lower in recipients of O-NT BM compared with those receiving young or O-Dox BM, indicating an age-related decline in reconstitution capacity that was prevented by early OSKM induction (Extended Data Fig. 3C, D). In tertiary transplantation, none of the recipients of O-NT BM survived, in contrast to mice transplanted with young or O-Dox cells (Extended Data Fig. 3E), suggesting that Dox treatment prevented age-associated HSC exhaustion.

To further confirm the rejuvenating effect of Dox on HSC function, we next performed competitive BM transplantation assays using O-NT and O-Dox donor cells (Fig. 2G). At all time points analyzed, donor chimerism in the peripheral blood was higher in recipients of O-Dox cells than in those receiving O-NT cells (Fig. 2H). This effect was particularly pronounced within the lymphoid compartment (Extended Data Fig. 3F–I). Notably, Dox also rescued the myeloid differentiation bias observed in O-NT donor-derived cells (Extended Data Fig. 3J). Consistent with these findings, BM analysis performed 4 months after transplantation revealed increased reconstitution of LSK cells, HSCs, and lymphoid-biased HSCs in recipients of O-Dox cells compared with those receiving O-NT cells (Fig. 2I–K).

Altogether, these results show that transient induction of OSKM factors early in life is sufficient to prevent, partially or fully, most age-associated alterations in hematopoiesis, including defects in HSC differentiation, myeloid bias, quiescence, and reconstitution capacity. It also reduces the accumulation of DNA damage in aged HSCs. Remarkably, these effects persisted for almost the entire lifespan of the animals despite complete disappearance of exogenous OSKM expression shortly after treatment, indicating that the protective effects of transient reprogramming are not mediated by sustained transcriptional activity but rather through the establishment of a durable epigenetic state.

### Partial transient reprogramming late in life rescues HSC aging

We next asked whether transient reprogramming initiated late in life could reverse age-associated alterations that had already been established. Remarkably, Old 4TF mice treated with Dox at 20 months of age and analyzed 4 months later (late Dox group, Fig. 2L) displayed peripheral blood lymphocyte and myeloid cell counts that were restored to levels comparable to those observed in young mice (Fig. 2M). In the BM, late-life reprogramming reduced the age-associated expansion of LT-HSCs, as well as of myeloid- and MK-biased HSC populations (Fig. 2N). It also reduced the accumulation of γH2AX foci in HSCs (Fig. 2O) and improved HSC replating capacity in methylcellulose cultures compared with O-NT HSCs (Extended Data Fig. 3K). Importantly, transient induction of OSKM late in life was sufficient to enhance the hematopoietic reconstitution potential of aged BM in competitive transplantation assays, with the most pronounced effect observed in the lymphoid compartment (Fig. 2P, Q and Extended Data Fig. 3L–O), while also reducing the age-associated myeloid differentiation bias (Extended Data Fig. 3P). BM analysis of recipients showed increased reconstitution of LSK cells, HSCs, and lymphoid-biased HSCs following transplantation with O-Dox late donor cells compared with O-NT cells, similar to the effects observed with O-Dox donor cells (Fig. 2R-T).

Taken together, these findings demonstrate that transient reprogramming can both delay hematopoietic aging and partially restore and the functional capacity of aged HSCs and their DNA integrity.

### Partial transient reprogramming early in life reduced age-associated transcriptomic changes

To determine whether the functional changes observed in aged HSCs following early-life reprogramming were associated with transcriptomic rejuvenation, we performed bulk transcriptomic analysis on sorted HSCs from young and old mice that had or had not been treated with Dox early in life. To assess whether Dox induction exerted an early effect on the transcriptome, an additional group was included in which HSCs were isolated one month after Dox treatment (Y-Dox mice, Fig. 3A). Unsupervised clustering of all differentially expressed genes (DEGs, FDR <0.1, [Log2FC] ≥0.4) showed that Y-NT and Y-Dox samples clustered together and that O-Dox samples are closer to these young samples than to the O-NT samples (Fig. 3B and Supplementary Table 1).

**Figure 3:**
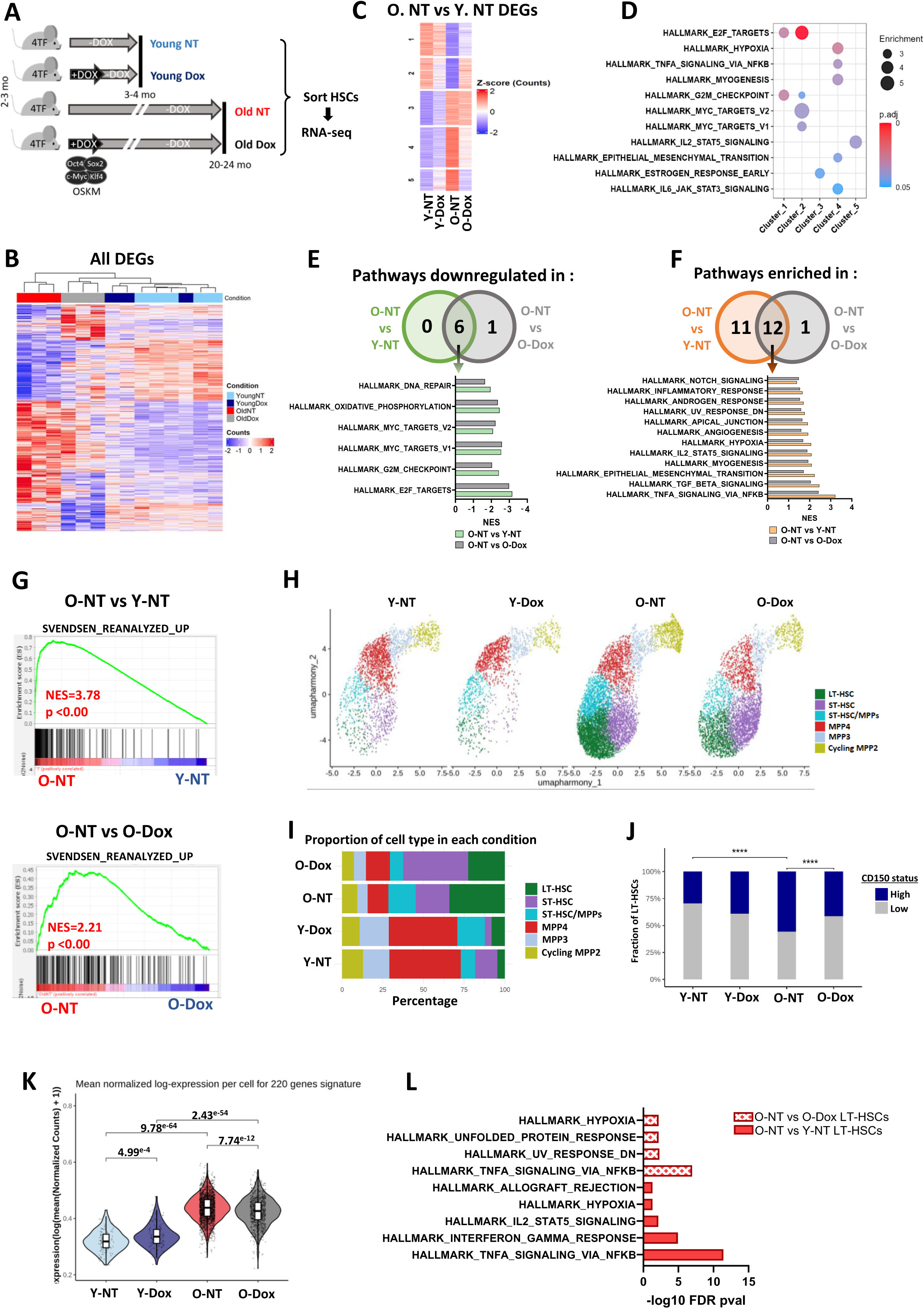
Partial reprogramming early in life prevents age-associated transcriptomic alterations. **(A-G)** Bulk RNA-Seq on HSCs (LSK CD34^-^ Flk2^-^) from Y-NT (n=5), Y-Dox (n=3), O-NT (n=3) and O-Dox (n=3) mice. (A) Experimental groups; (B) Clustering of samples based on normalized expression of DEGs (FDR < 0.1, log2FC ≥ 0.4) identified from all comparisons; (C) Heatmap of median expression in the 4 groups of the DEGs identified in the O-NT vs Y-NT comparison; (D) GSEA analysis showing HALLMARK gene sets significantly enriched (FDR <0.05) in clusters 1-5 in (C); (E, F) Venn diagram (up) and normalized enrichment scores (NES) (down) of HALLMARK gene sets commonly downregulated (FDR< 0.05) (E) or upregulated (F) in O-NT vs Y-NT (green or orange) and O-NT vs O-Dox (grey); (G) GSEA analysis performed with the published list of 220 aging DEGs from Svendsen et al.^31^ for the indicated comparisons. **(H-L)** Single cell RNA-Seq. LSK Flk2-cells from a pool of 6 Y-NT,6 Y-Dox, 3 O-NT and 3 O-Dox mice. (H) UMAP profile of cells and (I) their proportions among each condition; (J) Proportions of LT-HSC CD150^high^ and CD150^low^ among each condition. Adjusted p-values are reported as **** (p < 0.0001); (K) Aging signature score for each individual LT-HSC defined as the log-transformed mean normalized expression to of all genes in the signature set from Svendsen et al.^31^ Adjusted-pvalues are shown; (L) HALLMARK gene sets of upregulated DEGs (FDR <0.05; log2Foldchange > 0.25) in LT-HSCs for the indicated comparisons. J, K: Pairwise comparisons were performed using two-sided Wilcoxon rank-sum tests.

Aging induced 1036 DEGs as compared to Y-NT HSCs and 749 DEGs as compared to O-Dox HSCs, off which 60% and 53% where upregulated, respectively. By contrast, only few or no DEGs could be found between the Y-Dox control group and the O-Dox or Y-NT mice (Extended Data Fig. 4A), suggesting that Dox has no early effect on the transcriptome but rather largely prevented age-associated transcriptomic changes.

Heatmap clustering analysis of the DEGs in O-NT vs Y-NT HSCs in the 4 conditions further showed that the majority of these DEGs were fully or partially restored toward youthful expression levels in O-Dox HSCs (Fig. 3C). In agreement with previous reports^2, 10, 30–32^, age-upregulated DEGs (clusters 3–5) were enriched in pathways related to inflammation, cytokine signaling, hypoxia, and epithelial– mesenchymal transition (EMT), whereas age-downregulated DEGs (clusters 1–2) were enriched for cell cycle–associated genes (Fig. 3D).

To further evaluate the extent to which early partial reprogramming modulated age-associated transcriptomic programs and biological processes, we performed gene set enrichment analysis (GSEA). Specifically, we identified pathways enriched or downregulated in O-NT vs Y-NT that also remained similarly enriched or downregulated in O-NT vs O-Dox comparisons, thereby indicating rescue by OSKM induction. Among processes downregulated in O-NT in both comparisons, this analysis identified 6 HALLMARK and 118 REACTOME overlapping pathways (FDR < 0.05; Supplementary Table 2) primarily related to cell cycle regulation, DNA repair, and epigenetic processes (Fig. 3E and Extended Data Fig. 4B). Likewise, 12 HALLMARK and 4 REACTOME gene sets enriched in both O-NT vs Y-NT and O-Dox vs O-NT comparisons were associated with inflammatory responses, cytokine signaling, EMT, and TGFβ signaling (Fig. 3F, Extended Data Fig. 4C and Supplementary Table 2). The comparable normalized enrichment scores observed across these overlapping gene sets in the O-NT vs Y-NT and O-NT vs O-Dox comparisons indicate that the transcriptomic profile of O-Dox HSCs more closely resembles that of young HSCs.

To validate these findings, we examined an HSC aging signature reported by Svendsen et al. ^31^ following reanalysis of multiple public datasets. Of the 220 genes included in this signature, 52% overlapped with DEGs identified in the O-NT vs Y-NT comparison. Importantly, this aging signature was strongly enriched in both O-NT vs Y-NT and O-NT vs O-Dox comparisons (Fig. 3G). Finally, consistent with previous studies, gene set variation analysis (GSVA) revealed that HSC vs MPP4 lymphoid progenitor-^33^ and megakaryocyte (MK)-biased HSC signatures^34^ were markedly enriched in O-NT HSCs, whereas the high-output HSC signature was reduced (Extended Data Fig. 4D). Importantly, the magnitude of these age-associated changes was attenuated in O-Dox HSCs, linking transcriptomic remodeling to the functional phenotypes observed.

Single-cell RNA sequencing (scRNA-seq) analysis of the LSK Flk2⁻ cell population, which is enriched for HSCs, was performed across the four experimental groups. UMAP-based clustering confirmed the age-associated increase in the proportion of HSCs and CD150^hi^ cells, together with the reduced ST-HSC/LT-HSC ratio observed Old-NT mice, all of which were partially restored by early reprogramming (Fig. 3H– J). Notably, clustering analysis did not identify any HSC population specifically enriched in either the Y-Dox or Old-Dox groups, indicating that OSKM expression did not drive the emergence or expansion of a distinct HSC subset with a youthful transcriptional identity. Consistent with the bulk-RNA-Seq results, gene expression analysis at LT-HSC level showed that early reprogramming attenuated increased expression of the aging signature genes in LT-HSCs and of inflammatory and stress pathways that are commonly upregulated in O-NT LT-HSC compared to both Y-NT and O-Dox LT-HSCs (Fig. 3K-L, Extended Data Fig. 4E and Supplementary Table 3) ^31^

Collectively, these findings demonstrate that transient early-life induction of OSKM factors does not induce immediate significant transcriptomic changes in young mice but prevents the majority of age-associated transcriptomic alterations in HSCs.

### Partial transient reprogramming early in life reduced age-associated H3K9me3 loss at TEs in HSCs

The delay of age-associated transcriptomic alterations upon transient early-life reprogramming raised the possibility of a persistent epigenetic effect of OSKM induction. This prompted us to investigate the impact of Dox treatment on the epigenome. H3K9me3, a histone modification associated with constitutive heterochromatin, has been reported to decline globally with aging across multiple organisms and tissues^35^. Previous studies, including our own, have also demonstrated a marked reduction of this epigenetic mark in aged mouse and human HSCs^17, 36, 37^. We therefore examined whether transient early-life reprogramming could counteract the age-related decrease in H3K9me3 levels in HSCs. Immunofluorescence (IF) analyses confirmed a significant global reduction of H3K9me3 in HSCs from O-NT mice compared to Y-NT (Fig. 4A). Remarkably, early-life Dox treatment fully restored H3K9me3 levels to those observed in young HSCs. In contrast, Dox administration in WT mice failed to prevent the age-associated loss of H3K9me3 (Extended Data Fig. 5A), indicating that this effect specifically depends on OSKM induction.

**Figure 4:**
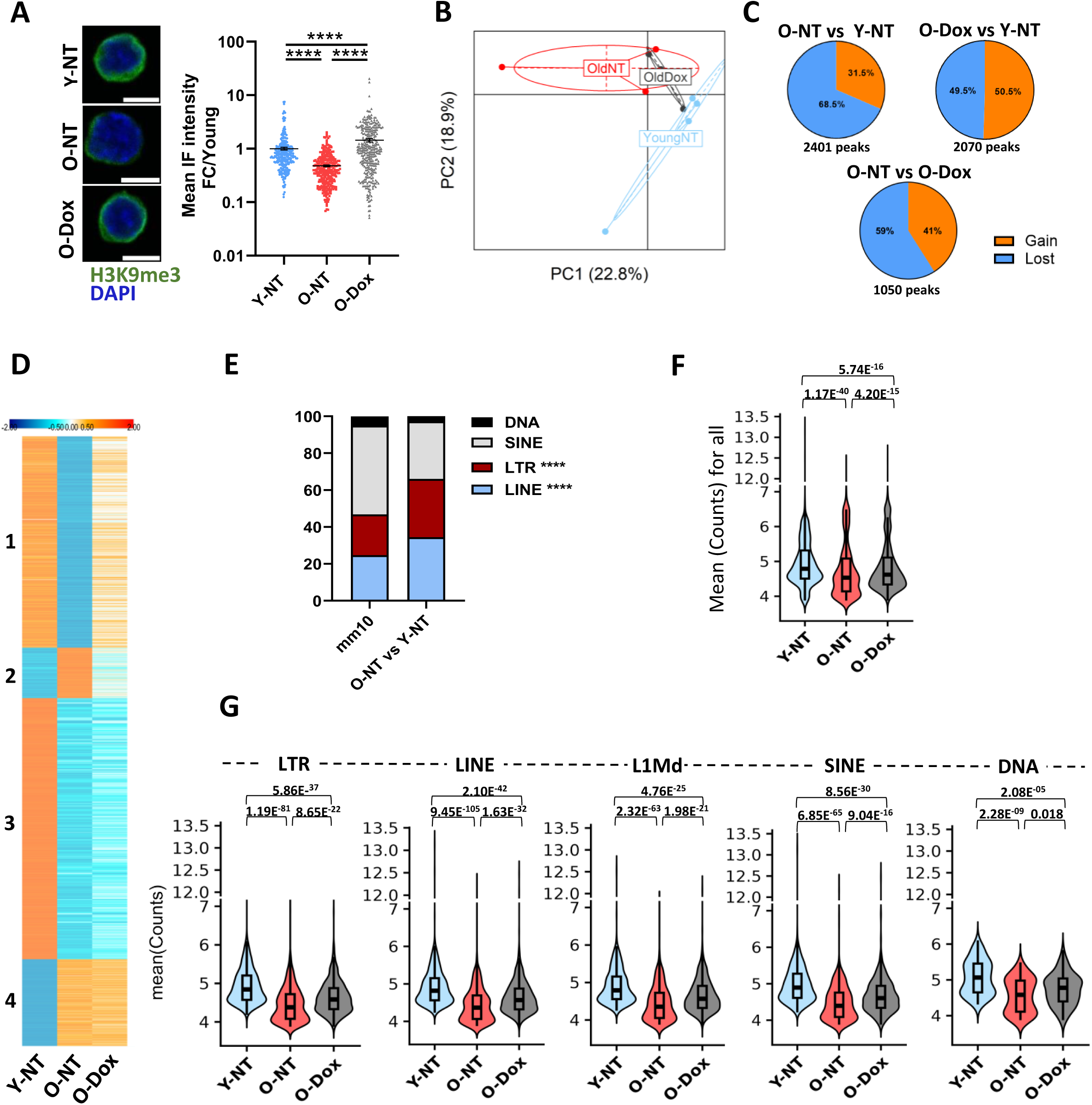
Partial reprogramming early in life limits the loss of H3K9me3 at TEs with age. **(A)** H3K9me3 IF representative images and intensity in HSCs. Y-NT (n =4), O-NT (n =5) and O-Dox (n=5) mice from 3 independent experiments. Each dot represents one HSC. Mean +/- SEM. One-way ANOVA with multiple comparison test. Adjusted p-values are reported as **** p<0.0001. Scale bars: 5µm. **(B-G)** H3K9me3 CUT&Tag. HSCs from Y-NT (n=4), O-NT (n=4) and O-Dox (n=4) mice. (B) PCA based on the top 500 most variable H3K9me3 peaks across samples; (C) Proportion of significantly lost (blue) and gained (orange) peaks for the indicated comparison (p-val <0.05); (D) Heatmap showing H3K9me3 concentration at TEs within O-NT vs Y-NT differential peaks, in the 3 groups; (E) Distribution of different TE families in the mouse mm10 genome (left) and among the O-NT vs Y-NT differential TEs (right). Hypergeometric test; (F, G) H3K9me3 concentration at all TEs (F) and at TEs from the different families (G) losing H3K9me3 with age retrieved in D. Statistical significance was assessed using Wilcoxon rank-sum tests, with adjusted p-values for multiple comparisons using the Holm method.

To further characterize these findings at the genome-wide level, we performed CUT&Tag analyses. Given that H3K9me3 is predominantly enriched at TEs in HSCs, sequencing data were analyzed using both uniquely and multiply mapped reads which were randomly assigned to one of their best genomic positions, as previously described^19^. Similar to the transcriptomic analyses, PCA revealed that O-Dox samples clustered between Young and O-NT groups (Fig. 4B). Differential peak analysis (p < 0.05) identified 2401, 2070, and 1050 peaks in the comparisons O-NT vs Young, O-Dox vs Young, and O-NT vs O-Dox, respectively (Fig. 4C; Supplementary Table 4). In all comparisons, differential peaks were predominantly located within intergenic and intronic regions (Extended Data Fig. 5B). Consistent with the IF data, most differential peaks in O-NT HSCs (68.5%) exhibited a loss of H3K9me3 relative to young cells. Notably, this proportion was reduced in HSCs from early reprogrammed O-Dox mice (Fig. 4C). Moreover, the comparable proportion of lost peaks observed in the O-NT vs Young and O-NT vs O-Dox comparisons suggests that Dox treatment effectively rescued the age-associated decline in H3K9me3 levels.

TE annotation of O-NT versus Young differential peaks revealed that a large proportion of TEs (4,406; 77%) were associated with differential peaks showing a loss of H3K9me3 during aging, compared with only 1,283 TEs (23%) associated with regions showing an age-related gain of this mark. Heatmap clustering analysis of H3K9me3 enrichment at these TEs showed that transient reprogramming partially prevented the age-related decline in H3K9me3 levels (Fig. 4D). TE class distribution further showed that LTRs and LINEs were significantly enriched among age-associated H3K9me3 differential peaks compared with their overall genomic representation (Fig. 4E). H3K9me3 enrichment at peaks bearing TEs was significantly reduced in O-NT HSCs relative to both Young and O-Dox groups (Fig. 4F). This preservation effect of Dox extended across all TE classes, as well as the evolutionarily recent mouse LINE-1 subfamily L1Md (Fig. 4G).

Collectively, these findings demonstrate that transient early-life reprogramming prevents the age-associated loss of H3K9me3 at TEs in HSCs.

### TE overexpression in aged HSCs promotes DNA damage and functional decline, an effect prevented by early reprogramming

To determine whether the age-associated loss of H3K9me3 at TEs, and its rescue by Dox treatment, were associated with TE expression, we first evaluated dsRNA accumulation using the J2 antibody, which specifically recognizes dsRNA species. IF analyses revealed a strong cytoplasmic dsRNA signal in HSCs from untreated Old mice, whereas this signal was reduced to levels comparable to young HSCs in O-Dox mice (Fig. 5A). Similar results were observed after late reprogramming in Old mice (Extended Data Fig. 6A). These findings suggest that TE expression increases with age and is prevented by transient reprogramming. Consistent with this observation, RNA-seq analysis demonstrated increased expression of all major TE classes in O-NT HSCs, which was markedly reduced following early Dox treatment (Fig. 5B, Extended Data Fig. 6B, Supplementary Table 5). Interestingly, early Dox treatment also reduced the age-associated upregulation of long (> 4 kb) L1Md elements (Fig. 5B), which are more likely to retain intact coding sequences for LINE-1 endonuclease and reverse transcriptase activities and which contribute to irradiation- and chronic inflammation-induced DNA damage and dysfunction in HSCs^15, 19, 20^. We therefore investigated whether increased L1Md expression during aging contributes to the elevated DNA damage and reduced regenerative capacity observed in aged HSCs. To this end, we used lamivudine (3TC), a reverse transcriptase inhibitor previously shown to inhibit LINE-1 reverse transcriptase activity^15, 20^. In vitro treatment of O-NT HSCs with lamivudine restored their clonogenic potential during serial replating assays at passages 2 and 3 (Fig. 5C, D). In contrast, lamivudine treatment had no significant effect on the replating activity of HSCs isolated from O-Dox mice reprogrammed with transient OSKM induction, either early or late in life (Fig. 5D and Extended Data Fig. 6C). These results suggest that age-associated L1Md activation and the resulting DNA damage contribute substantially to the loss of HSC self-renewal capacity, and that transient reprogramming limits these deleterious effects to preserve HSC function.

**Figure 5:**
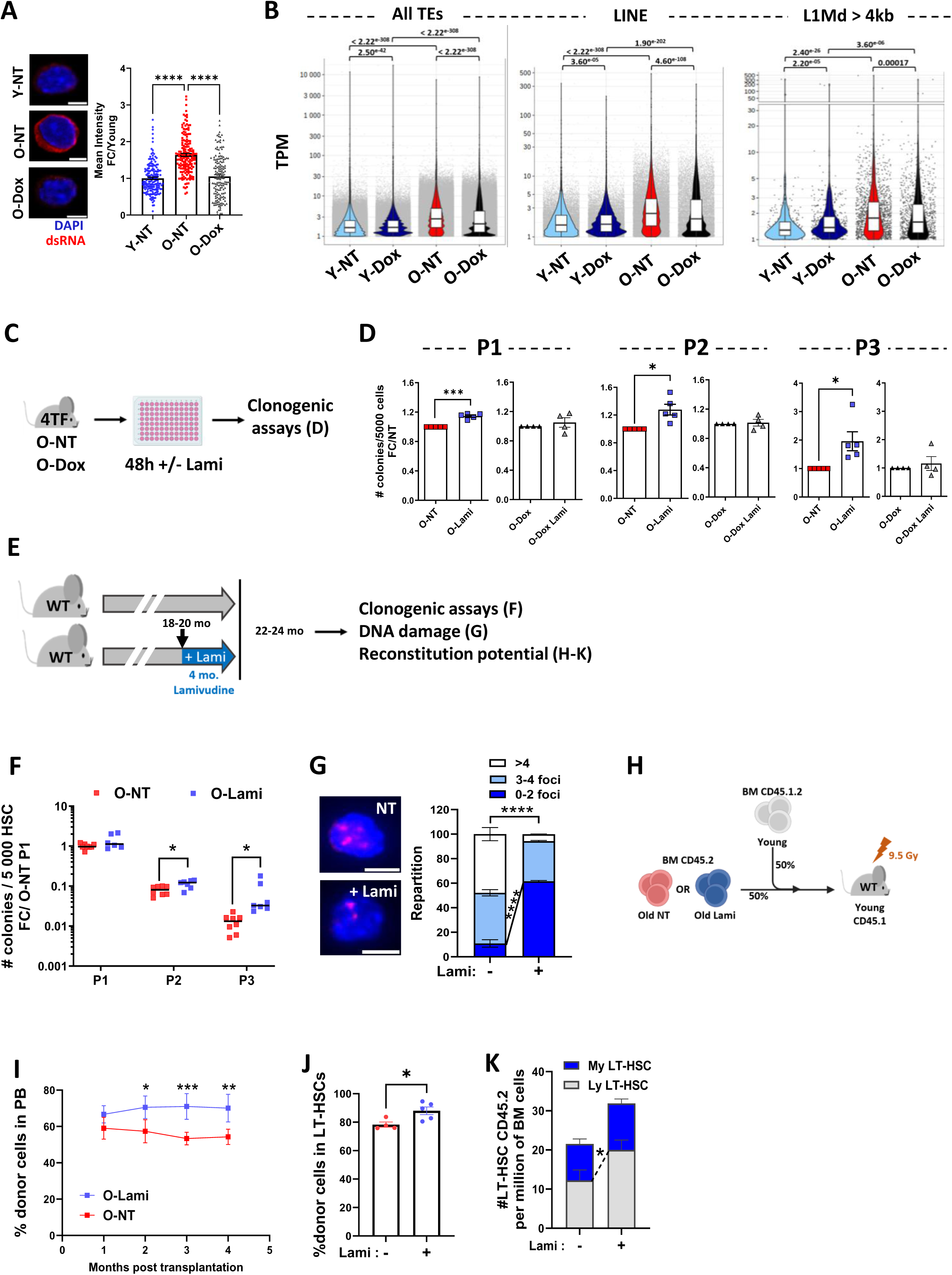
Partial reprogramming early in life prevents TE overexpression with age to mitigate HSC functional decline. **(A)** dsRNA IF representative images and intensity. HSCs from Y-NT (n=3), O-NT (n=3) and O-Dox (n=3) mice from 2 independent experiments. Results from at least 45 HSCs per mouse. Each dot represents one HSC. Means +/- SEM. One-way ANOVA with multiple comparisons test, adjusted p-values are reported as **** p<0.0001. Scale bars on the picture represent 5µm. **(B)** Violin plots showing the expression of TEs copies (All TEs, LINEs and L1Md > 4 kb) quantified using the SQUIRE pipeline into the RNA-Seq data set (Fig. 3A). Each TE copy compute mean TPM value filtered by threshold TPM >=1 in at least one condition are shown. One-way ANOVA with multiple comparisons test. Adjusted p-values are shown. **(C, D)** Experimental design (C) and number of colonies (D) generated from 5000 HSCs from O-NT and O-Dox mice treated of not with of lamivudine (10 µM) for 48h at each plating. Each dot represents an individual mouse. 2 independent experiments. Results are normalized to the non-treated group. Paired t-test, * p-val <0.05, *** p-val <0.001. **(E-K)** Effect of lamivudine (Lami) treatment of aged WT mice *in vivo*. (E) Experimental design; (F) Number of colonies generated at platings 1, 2, 3 by 5000 HSCs from mice either non-treated (NT) or treated with lami for 4 months (O-Lami). Each dot represents an individual mouse. 2 independent experiments. Means +/- SEM. Multiple t tests, * p-val <0.05; (G) γH2AX foci representative images and repartition. At least 50 HSCs per mouse treated (n=2) on not (n=3) with lamivudine. Scale bars represent 5µm in pictures; (H-K) Competitive BM transplantation between Old +/- Lami BM cells and Young BM cells into lethally irradiated young mice. (H) experimental design; (I) Percentage of donor-derived cells in peripheral blood over time. (J) Percentages of donor-derived cells into LT-HSCs in the BM of recipients and (K) numbers of My- and Ly-biased donor-derived LT-HSCs, 4 months post-transplantation. O-NT (n=4), O-Lami (n=5). Means +/- SEM. G, I, K: Two-way ANOVA with multiple comparisons test. Adjusted p-values defined as: * p<0.05; ** p<0.01; *** p<0.001; **** p<0.0001. J: t-test * p<0.05

To further assess the contribution of TEs to HSC aging in vivo, WT 20-month-old mice were continuously treated with lamivudine for four months, after which blood and BM parameters, HSC function and DNA damage were evaluated (Fig. 5E). After lamivudine treatment the age-associated myeloid bias observed in peripheral blood was slightly decreased whereas no effect was observed has on HSC numbers in the BM or HSC myeloid bias (Extended Data Fig. 6D, E). Consistent with our in vitro findings, HSCs isolated from lamivudine-treated mice displayed enhanced replating capacity (Fig. 5F) together with reduced DNA damage (Fig. 5G). Competitive bone marrow transplantation assays further demonstrated that lamivudine improved hematopoietic reconstitution capacity, enhanced HSC self-renewal, and reduced myeloid skewing in aged HSCs (Fig. 5H–K).

Altogether, these findings demonstrate that age-associated TE derepression contributes to increased DNA damage, HSC dysfunction, and myeloid bias during aging. They further suggest that transient early-life reprogramming preserves HSC function by preventing TE activation and its deleterious consequences.

### Early induction of OSKM factors prevents AP-1–associated chromatin opening and induction of inflammatory / EMT gene signature with age

Because OSKM factor expression in HSCs is transient, the above results suggest that the effects of reprogramming are stably inherited in aged HSCs, further supporting the notion that aging is largely driven by epigenetic mechanisms. To investigate further how transient expression of OSKM TFs^38^ could exert long-term effects on HSC function, we performed Assay for Transposase-Accessible Chromatin sequencing (ATAC-seq) on HSCs isolated from the four experimental groups: Y-NT, Y-Dox, O-NT, and O-Dox mice (Fig. 3A).

A clustering heatmap based on all differentially accessible regions (DARs; FDR < 0.1) showed that, similarly to the transcriptomic analyses, O-Dox samples clustered more closely with Y-NT and Y-Dox samples than with O-NT samples (Fig. 6A and Supplementary Table 6). No significant DARs were identified between the two young groups (Extended Data Fig. 7A), indicating that Dox treatment does not induce detectable chromatin remodeling in young HSCs. In contrast, 498 DARs were detected between O-NT and Y-NT HSCs, the majority of which (79%) exhibited increased accessibility (Extended Data Fig. 7A), consistent with the widespread chromatin opening previously reported in aged HSCs^8^. By comparison, among the 904 DARs identified between O-Dox and Y-Dox HSCs, most (81%) displayed reduced chromatin accessibility. Furthermore, chromatin regions in O-NT HSCs remained predominantly more accessible (99% of DARs) when compared with O-Dox HSCs, suggesting that Dox treatment prevented age-associated chromatin opening and may actively contribute to chromatin closing. Genomic annotation of DARs revealed a predominant localization within intergenic and intronic regions across all comparisons (Extended Data Fig. 7B).

**Figure 6:**
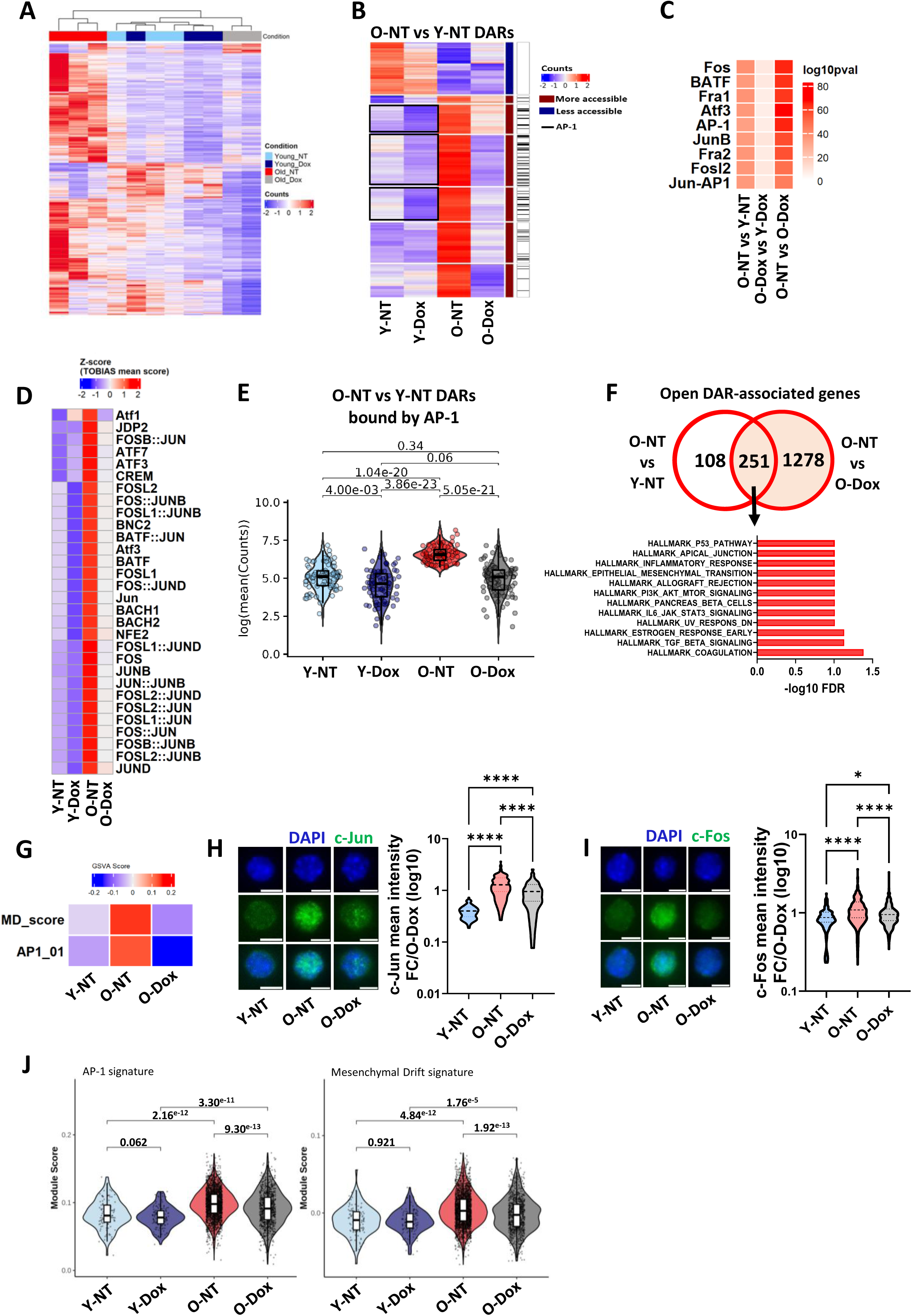
Partial reprogramming early in life prevents chromatin opening at AP-1 binding sites. **(A-E)** ATAC-seq of HSCs (LSK CD34^-^ Flk2^-^) from Y-NT (n=3), Y-Dox (n=3), O-NT (n=3) and O-Dox (n=2) mice. (A) Clustering of samples based on accessibility of DARs (FDR <0.1) identified from all comparisons; (B) Heatmap showing the median read counts of O-NT vs Y-NT DARs in all groups. The sidebar indicates DARs containing at least one JUN or FOS (AP-1) TF expected to be bound, as detected by TOBIAS analysis; (C) Heatmap showing the enrichment of the top 30 HOMER TF binding motifs (p ≤10^-30^) in O-NT vs Y-NT open DARs, compared to the non-significant background, in all indicated comparisons; (D) Heatmap showing the top 30 of most variable TF activities between O-NT vs Y-NT inferred from ATAC-seq footprinting using TOBIAS across Y-NT, Y-Dox, O-NT and O-Dox groups; (E) Violin plot representation of ATAC-seq signal at FOS and JUN footprint regions in O-NT vs Y-NT. Statistical significance was assessed using Wilcoxon rank-sum tests, with p-values adjusted for multiple comparisons using the Holm method. Adjusted p-values are shown. **(F)** Venn diagram and associated HALLMARK gene sets for common genes falling nearest to the open O-NT vs Y-NT and O-NT vs O-Dox DARs. **(G)** Bulk RNA-seq GSVA analysis showing the median score for AP-1 and mesenchymal drift (MD) score pathways. **(H, I)** IF representative images and intensity of c-Jun (H) and c-Fos (I) protein expression in HSCs from Y-NT (n=2), O-NT (n=3), O-Dox (n=3) mice. At least 50 cells per mouse. Violin plots presenting median and quartiles. Results are normalized on the O-Dox group. One-way ANOVA, adjusted p-values: * p<0.05; **** p<0.0001. **(J)** Expression levels of AP-1 and MD signatures in the LT-HSC population from single-cell RNA-Seq data. Pairwise comparisons were performed using two-sided Wilcoxon rank-sum tests with FDR correction. Adjusted p-values are shown.

Analysis of the median accessibility of age-associated peaks across experimental conditions demonstrated that a significant fraction of chromatin regions that became accessible in O-NT HSCs reverted toward youthful accessibility levels in O-Dox HSCs (Fig. 6B and Extended Data Fig. 7C). In contrast, the effect of Dox treatment on age-associated chromatin closing appeared more limited (Fig. 6B). TF motif analysis using HOMER detected no significantly enriched motifs (p < 10^-3^) among these closing DARs (Supplementary Table 7), likely because relatively few regions lost accessibility with age. TF motif analysis on peaks gaining accessibility in O-NT HSCs revealed strong enrichment AP-1 complex-forming TFs, including FOS, JUN, ATF and Fra when compared to both Y-NT and O-Dox HSCs (Supplementary Table 7). This is consistent with a recent report describing pervasive AP-1-associated chromatin opening across multiple aged tissues^38^. Among age-associated opening DARs, 79 out of 393 regions (20%) contained at least one predicted binding site for AP-1 FOS or JUN family members (Fig. 6B, right column). Importantly, while AP-1 binding site enrichment at DARs was low in the O-Dox vs Y-Dox comparison it remained strongly enriched in O-NT vs O-Dox (Fig. 6C). TF activity inferred from ATAC-seq footprinting using TOBIAS analysis^39^ further confirmed a genome-wide increase in AP-1 binding score in O-NT HSCs as compared to both Y-NT and O-Dox HSCs (Extended Data Fig. 7D and Supplementary Table 8), indicating that Dox treatment partially prevented age-associated activation of AP-1 TFs. To refine this analysis, TOBIAS scores were calculated for age-associated opening DARs after filtering RNA-seq data to retain only expressed TFs (see Methods). AP-1 family activity ranked among the strongest signals detected in O-NT open DARs and was markedly reduced in the O-Dox group (Fig. 6D). Consistent with these findings, DAR read counts at AP-1 binding sites identified within O-NT vs Y-NT DARs were significantly increased in O-NT HSCs and reduced in O-Dox HSCs, returning toward youthful levels (Fig. 6E).

AP-1 activity has been identified as a major barrier to reprogramming, and one of the earliest events during OSKM-mediated iPSC reprogramming is the closing of chromatin at AP-1–bound regulatory regions^40–43^. Interestingly, despite the absence of statistically significant DARs between Y-NT and Y-Dox HSCs, 77% of age-associated opening DARs containing at least one predicted AP-1 footprint (FOS or JUN) displayed reduced chromatin accessibility in the Y-Dox group compared to Y-NT (Fig. 6B, black boxes). Supporting these observations, Y-Dox HSCs exhibited a marked reduction in TOBIAS scores for AP-1 TFs (Fig. 6D). In addition, read counts at AP-1 binding sites identified within O-NT vs Y-NT DARs were significantly decreased in Y-Dox HSCs compared with Y-NT controls (Fig. 6E). These findings suggest that an early consequence of transient OSKM induction in HSCs is the selective closing of chromatin at AP-1-bound regulatory regions, and that this chromatin state is stably inherited during aging.

Previous studies have shown that AP-1 suppresses the mesenchymal-to-epithelial transition (MET), a process required for reprogramming, by activating an EMT transcriptional program^44^. Analysis of genes lying closest to the DARs identified 359 genes in the O-NT vs Y-NT comparison and 1,529 genes in the O-NT vs O-Dox comparison (Fig. 6F, Supplementary Table 6). Among these, 251 genes were common to both datasets, representing genes rescued by Dox treatment. These genes were enriched for EMT and pathways linked to EMT such as apical junction, TGFβ, and inflammatory/cytokine signaling pathways (Fig. 6F), consistent with increased AP-1 activity during aging. Similarly, genes associated with chromatin regions losing accessibility in Y-Dox vs Y-NT were enriched for the same transcriptional programs (Extended Data Fig. 8A).

Integration of RNA-seq and ATAC-seq data revealed that DEGs associated with DARs gaining chromatin accessibility in O-NT relative to either Y-NT or O-Dox HSCs were modestly but significantly upregulated in O-NT HSCs across both comparisons (Extended Data Fig. 8B). These DEGs were predominantly enriched for EMT and EMT-linked pathways, as well as additional AP-1-associated transcriptional programs, including inflammatory response, MAPK signaling, and UV response (Extended Data Fig. 8C).

Consistent with these observations, O-NT HSCs displayed increased expression of an aging-associated mesenchymal drift signature⁴⁵ together with a robust AP-1 transcriptional program, both of which were largely reversed following transient OSKM induction in O-Dox HSCs (Fig. 6G). In agreement with these transcriptional changes, protein levels of the AP-1 family members c-Jun and c-Fos were increased in O-NT HSCs, compared to both Y-NT and O-Dox HSCs (Fig. 6H, I), further corroborating the suppression of this pathway upon partial reprogramming.

To further assess AP-1 activity at the single-cell level, we analyzed AP-1 and mesenchymal drift pathway activity in LT-HSCs from scRNA-seq data. This showed that transient OSKM expression attenuated the activation of both AP-1 and EMT programs in LT-HSCs (Fig. 6J). These findings further support the conclusion that the rejuvenating effects of transient OSKM induction are impacting the LT-HSC compartment and are associated with suppression of AP-1-dependent transcriptional activity.

Collectively, these findings demonstrate that transient OSKM induction establishes a durable epigenetic memory in HSCs by selectively preventing age-associated chromatin opening at AP-1 regulatory elements. This chromatin remodeling is accompanied by repression of AP-1-dependent transcriptional programs, including EMT, inflammatory, and MAPK pathways associated with hematopoietic aging, thereby preserving a more youthful chromatin landscape and transcriptional state in aged HSCs.

### Early induction of OSKM factors prevents age-associated chromatin opening at TEs enriched for AP-1 binding sites

Analysis of ATAC-seq data at TE genomic loci revealed increased chromatin accessibility at 52 and 75 TEs, and decreased accessibility at 10 and 1 TEs, in O-NT HSCs compared with Y-NT or O-Dox HSCs, respectively (FDR < 0.1; Fig. 7A, Supplementary Table 9). In contrast, aging under Dox-treated conditions resulted predominantly in decreased accessibility, affecting 127 TEs, while only 34 TEs showed increased accessibility (Fig. 7A). Heatmap analysis of differentially accessible TEs in O-NT vs Y-NT demonstrated that Dox treatment reversed most of the age-associated chromatin opening events, whereas it had little effect on TEs that lost accessibility with age (Fig. 7B).

**Figure 7:**
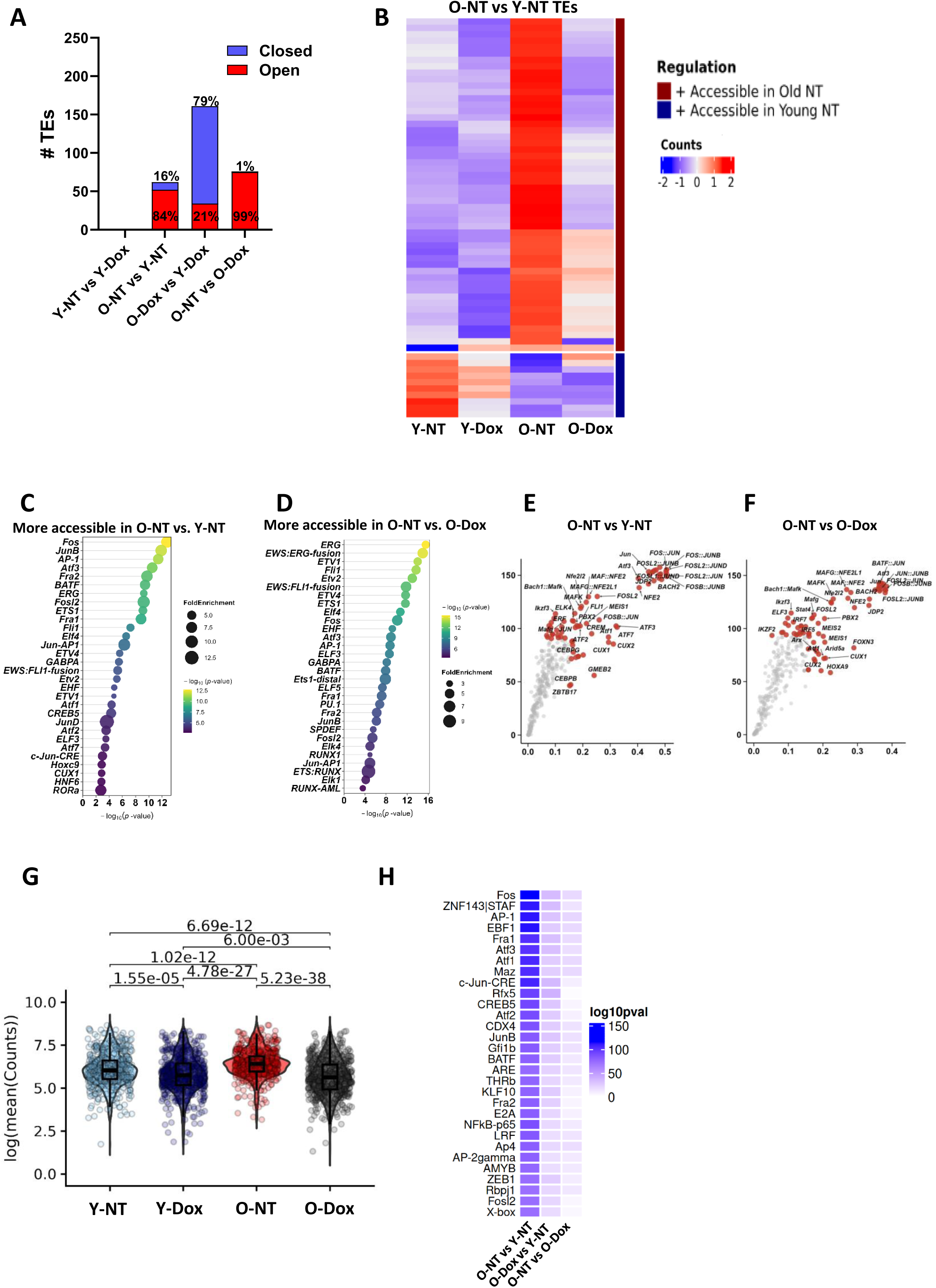
Partial reprogramming early in life prevents chromatin opening at TEs containing AP-1 binding sites. **(A-G)** ATAC-seq analysis at genomic TEs. (A) Number of open (red) or closed (blue) TEs (FDR <0.1) in the indicated comparisons; (B) Heatmap showing the mean read counts for differentially accessible TEs identified between O-NT vs Y-NT; (C, D) Top 30 HOMER motifs found at TEs more accessible in O-NT vs Y-NT (C) and O-NT vs O-Dox (D); (E, F) Volcano plots showing differential TF binding activity at TEs, as detected by TOBIAS, in O-NT vs Y-NT (E) and O-NT vs O-Dox (F) comparisons. Each point corresponds to a predicted TF. TF are ranked according to their binding score; (G) Violin plot showing the log2-transformed read counts at TE-associated peaks predicted to be bound by AP-1, obtained by TOBIAS footprinting intersecting all O-NT vs Y-NT peaks containing TEs. Statistical significance was assessed using Wilcoxon rank-sum tests, with p-values adjusted for multiple comparisons using the Holm method. **(H)** Heatmap showing the enrichment of the top 30 HOMER motifs (p ≤ 1e-05) found at TEs from CUT&Tag peaks significantly loosing H3K9me3 between O-NT vs Y-NT compared to non-significant background, and shown for all indicated comparisons.

TF motif analysis using HOMER (Supplementary Table 10) revealed strong enrichment for AP-1 family members, as well as ETS family TFs including ERG, Fli1 and RUNX1, at TEs gaining accessibility in O-NT cells across all comparisons (Fig. 7C, D). Furthermore, strong AP-1 activity at TE-associated peaks in O-NT HSCs, compared to both Y-NT and to O-Dox HSCs was confirmed by footprint analysis (Fig. 7E, F; Supplementary Table 11). Read counts at TE-associated peaks predicted to be bound by AP-1 were significantly increased in O-NT relative to both Y-NT and O-Dox HSCs (Fig. 7G). Dox treatment also induced an early reduction in chromatin accessibility at these sites in Y-Dox HSCs. Finally, HOMER analysis of TEs losing H3K9me3 with age identified AP-1 binding motifs among the most enriched TF binding sites (Fig. 7H; supplementary Table 12). This enrichment was markedly reduced in O-Dox HSCs, supporting the conclusion that transient OSKM induction triggered epigenetic silencing at AP-1-bound TEs.

### Transient OSKM induction prevents stress-induced HSC loss of function

AP-1 TFs are activated downstream of multiple stress-response pathways, including cytokine signaling, inflammatory stimuli, and DNA damage^45–47^. We therefore investigated whether transient OSKM induction could mitigate stress-induced HSC dysfunction associated with TE expression. To address this question, we used a premature aging model induced by chronic inflammation induced by LPS in which TEs have previously been shown to contribute to DNA damage accumulation and HSC functional decline^20^.

To determine whether transient OSKM induction could preserve HSC function under inflammatory stress, 4TF mice were treated with repeated low doses of LPS for one month, either beginning one month after Dox treatment or without prior Dox exposure, and HSC reconstitution activity was subsequently assessed by competitive BM transplantation assays (Fig. 8A). Dox treatment alone did not significantly affect hematopoietic reconstitution in the absence of LPS (Extended Data Fig. 9A). In contrast, transient OSKM induction markedly improved the reconstitution potential of cells from LPS-treated mice, as evidenced by increased donor chimerism in peripheral blood (Fig. 8B). This improvement was particularly pronounced in the lymphoid compartment, whereas the effect was more modest in myeloid cells (Extended Data Fig. 9B, C). Dox treatment also partially mitigated the LPS-induced reduction in HSPC and HSC reconstitution in the BM (Fig. 8C, D).

**Figure 8:**
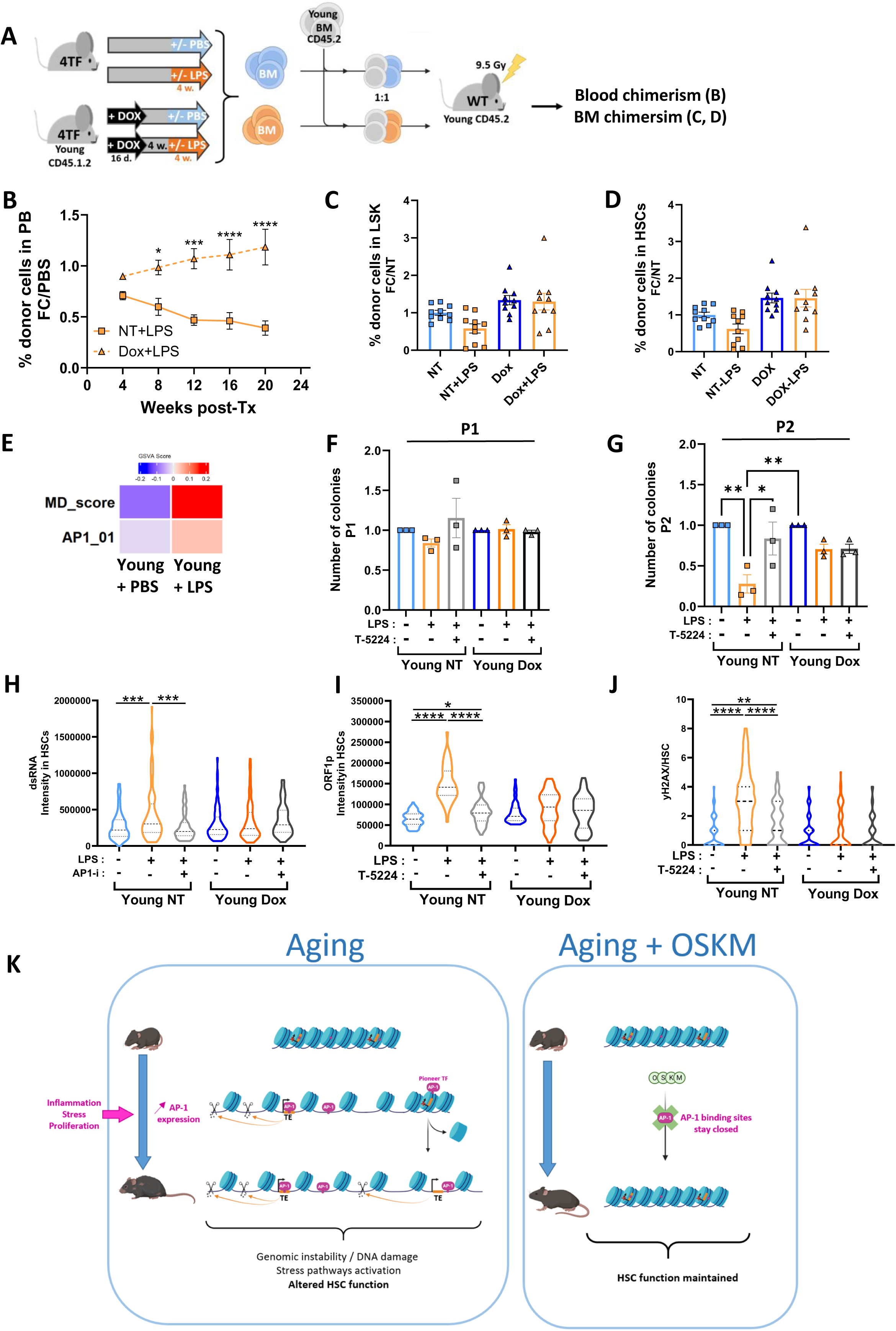
Partial reprogramming prevents chronic LPS-induced HSC loss of function. **(A)** Experimental design for competitive transplantations with BM cells from Young 4TF mice treated or not with Dox and submitted or not or to chronic LPS treatment during 1 month. **(B)** Percentages of donor-derived cells in the peripheral blood of recipients. Results are shown as the fold-change of LPS over PBS control group. Means +/- SEM, n=10 mice per group from 2 independent experiments. **(C, D)** Percentages of donor-derived cells into LSK cells (C) and HSCs (D) of recipient 5 months after transplantation. Each dot represents a mouse. Means +/- SEM. **(E)** GSVA analysis of the AP-1 and MD pathways in published RNA-seq data of HSCs from mice subjected to chronic LPS treatment for one month ^20^. **(F, G)** Number of colonies generated from HSCs cultured during 7 days in methylcellulose in the presence or absence of LPS (10 µg/mL) and the AP-1 inhibitor, T-5224 (10 µM), at passages 1 (F) and 2 (G) after replating in the absence of LPS or inhibitor. Results are normalized to the data obtained in the non-treated condition for each group. Each dot represents a mouse. Means +/- SEM. **(H-J)** dsRNA (H), L1 ORF1p (I) and γH2Ax foci (J) quantified by IF in HSCs after 48 hours of treatment with or without LPS (10 µg/mL) and T-5224 (10 µM) in vitro. Violin plots presenting median and quartiles. Independent pools of 3 mice per group. **(K)** Model suggesting that partial reprogramming early in life preserves chromatin homeostasis during HSC aging and upon stress, by avoiding chromatin remodeling at AP-1 binding sites and TE derepression, thereby maintaining a more youthful chromatin landscape, DNA integrity and HSC function. B: Two-way ANOVA with multiple comparisons test; C, D, F, G, H, I, J: One-way ANOVA with multiple comparisons test. Adjusted p-values are defined as: * p<0.05; ** p<0.01; *** p<0.001; **** p<0.0001

GSVA analysis of our previously published RNA-seq data of HSCs sorted after one month LPS treatment^20^ identified increased in both AP-1 and mesenchymal drift signatures as compared to control mice that received PBS (Fig. 8E). We therefore investigated the contribution of AP-1 signaling to reprogrammed HSC responses to LPS stimulation. Purified HSCs were seeded in methylcellulose in the presence or not of LPS, AP-1 inhibitor (T-5224), or the combination of both. After 7 days of primary culture, no significant differences in colony numbers were observed between the different conditions (Fig. 8F). However, following replating in fresh methylcellulose in the absence of LPS or AP-1 inhibitor, LPS-treated HSCs from 4TF NT mice exhibited a marked reduction in colony-forming capacity compared with PBS-treated controls, indicative of impaired self-renewal activity (Fig. 8G). Importantly, this defect was prevented by the presence of AP-1 inhibition during the primary culture period. In contrast, neither LPS nor AP-1 inhibition affected the colony-forming capacity of HSCs isolated from Dox-treated mice (Fig. 8F, G). Moreover, AP-1 inhibition alone had no effect on the colony-forming ability of either NT or Dox HSCs in the absence of LPS stimulation (Extended Data Fig. 9D).

LPS treatment in vitro for 48h was sufficient to induce the accumulation of dsRNA, LINE-1 ORF1 proteins (ORF1p) and γH2AX foci in NT HSCs whereas HSCs isolated from Dox-treated mice remained largely protected from this response (Fig. 8H-J and Extended Data Fig. 9E-G). These findings indicate that LPS acts directly on HSCs to promote L1 expression and that transient OSKM induction confers resistance to this effect. Notably, co-treatment with the AP-1 inhibitor prevented LPS-induced both dsRNA, ORF1p and γH2AX foci accumulation in NT HSCs (Fig. 8H-J and Extended Fig. 9E-G). Consistent with the clonogenic data, AP-1 inhibition had no detectable effect in Dox-treated HSCs, suggesting that prior transient OSKM induction in vivo has blocked HSC ability to trigger AP-1-dependent TE expression and TE-induced DNA damage in response to LPS.

Together, these results demonstrate that transient OSKM induction does not intrinsically enhance HSC function under steady-state conditions but instead protects HSCs from inflammation-induced functional decline. Our findings support a model in which transient reprogramming enhances HSC resilience to inflammatory stress, at least in part through the attenuation of AP-1-dependent TE activation and the consequent reduction in TE-associated stress responses and DNA damage pathways.

## Discussion

Over the last decade, epigenetic alterations have emerged as major candidates to explain the persistent dysfunction of aged HSCs transplanted in young environment^4^. Old HSCs exhibit widespread changes in DNA methylation, chromatin accessibility, histone modifications, transcriptional regulation, and higher-order chromatin organization, suggesting that aging results from the progressive accumulation of stable, yet potentially reversible, molecular alterations that become durably encoded over time. Here, we show that a brief transient induction of the Yamanaka factors is sufficient to durably remodel the trajectory of HSC aging. A single pulse of OSKM expression early in life attenuated many of the molecular and functional hallmarks of physiological aging, including myeloid skewing, loss of regenerative capacity, heterochromatin erosion, TE activation, DNA damage accumulation, and AP-1-associated chromatin remodeling. Remarkably, these effects persisted throughout most of the lifespan despite the complete disappearance of exogenous OSKM expression shortly after treatment, indicating that the protective effects of transient reprogramming are not mediated by sustained transcriptional activity, but rather by the establishment of a durable molecular state that slows the acquisition of age-associated alterations in HSCs.

Several interventions have been reported to delay some aspects of HSC aging to varying degrees^48–50^. Our experiments indicate that transient OSKM induction late in life remains capable of reversing several phenotypic and functional hallmarks of HSC aging, long after their establishment. These observations suggest that OSKM does not merely prevent the acquisition of age-associated alterations but can also reverse at least part of an existing aged state, supporting the notion that key aspects of HSC aging remain reversible. This is consistent with previous studies demonstrating old HSC rejuvenation by modulation of mTOR signaling, restoration of polarity through Cdc42 inhibition or lowering nuclear envelope tension through RhoA inhibition^7, 37, 51^. Integrated scRNA-seq analyses comparing Y-NT, Y-Dox, O-NT, and O-Dox HSCs failed to identify specific HSC populations that emerged early in life and were subsequently maintained with age in Dox-treated mice. Together, these findings suggest that rejuvenation results from the remodeling of aging-associated chromatin states across the entire HSC compartment, rather than from the emergence, expansion, or persistence of a specific HSC subset with a younger state identity.

Among the molecular alterations induced by transient reprogramming, AP-1-associated chromatin remodeling emerged as one of the earliest and most prominent features. AP-1 TFs are pioneer factors capable of remodeling the chromatin that occupy a central position in cellular stress responses downstream of inflammatory cytokines, MAPK signaling, oxidative stress, DNA damage, and diverse environmental insults^45–47, 52^. Although the mechanisms through which transiently expressed OSKM factors induce rejuvenation remain incompletely understood, AP-1 family members are well-established barriers to somatic cell reprogramming. Studies of iPSC reprogramming have demonstrated that AP-1-bound regions are among the earliest genomic loci to undergo chromatin closure following OSKM induction and that persistent AP-1 activity antagonizes, whereas its inhibition facilitates, acquisition of cellular plasticity^40–43^. Recent multi-tissue analyses have further identified increased AP-1 activity as a conserved hallmark of mammalian aging, associated with widespread chromatin opening and activation of inflammatory programs^38^. Our findings strongly support the idea that these observations are mechanistically linked. We found that genomic regions displaying age-associated increases in accessibility were highly enriched for AP-1 motifs, whereas transient OSKM induction markedly limited this process. Notably, although transient OSKM expression induced minimal transcriptional and chromatin remodeling in young HSCs, it was associated with an early reduction in accessibility at AP-1-bound regions, long before measurable aging phenotypes emerged. This observation suggests that AP-1-associated chromatin remodeling represents an initiating event rather than a secondary consequence of rejuvenation. AP-1 family members inhibit somatic cell reprogramming by promoting EMT through activation of mesenchymal transcriptional networks. Interestingly, recent studies have proposed that aging itself is accompanied by a progressive "mesenchymal drift" across multiple tissues^53^. Consistent with these observations, aged HSCs displayed increased c-Jun and c-Fos protein expression together with enrichment of mesenchymal drift, inflammatory and AP-1-associated EMT signatures, all of which were attenuated by transient reprogramming. Although HSCs are not epithelial cells and do not undergo EMT in the classical sense, these findings suggest that one of the earliest consequences of transient OSKM reprogramming may be the establishment of a long-lasting epigenetic state that is refractory to stress-induced AP-1 activation (Fig. 8 K).

The prominence of AP-1 signatures in our datasets is particularly intriguing in light of recent studies describing long-term inflammatory memory in different types of cells. In epidermal and intestinal stem cells, transient inflammatory stimuli induce persistent chromatin accessibility changes at AP-1 regulatory regions that facilitate future responses to tissue injury^54, 55^. A similar mechanism has been identified in memory T cells^52^. Epigenetic memory of inflammatory challenges has also been reported in HSCs^56, 57^. Most notably, recent studies identified inflammatory memory HSCs displaying persistent AP-1-associated transcriptional programs that accumulate with age in humans and are enriched in clonal hematopoiesis^58, 59^. Conversely, TET2-mutant HSPCs exhibit intrinsic epigenetic silencing of AP-1 TFs and attenuated transcriptional responses to inflammatory stimuli, a feature thought to underlie their resilience and selective expansion under chronic inflammatory conditions^60^. Together, these observations raise the possibility that AP-1 functions not only as an immediate stress-responsive TF but also as a key mediator through which environmental insults become durably encoded within the HSC epigenome and involved in HSC aging. Our findings strongly support this model. Several studies from our group and others have shown that the same molecular hallmarks—including chromatin opening, heterochromatin erosion, TE derepression, and DNA damage accumulation—recurrently emerge across physiological aging and irradiation- or chronic-inflammation-induced premature aging^8, 10, 15–20^. In the present study, transient OSKM reprogramming in young mice rescued the reconstitution potential of both aged and chronic LPS-treated HSCs. Moreover, pharmacological inhibition of AP-1 prevented the LPS-mediated reduction in clonogenic potential in vitro, while exerting no additive effect in Dox-treated HSCs, either in the presence or absence of LPS. These findings suggest that a major function of transient OSKM reprogramming is to prevent the establishment of stress-induced AP-1 programs rather than simply counteracting their downstream consequences (Fig. 8K).

Finally, our data establish TE dysregulation as an important contributor to physiological HSC aging. We and others previously demonstrated that irradiation, chemotherapy, and chronic inflammatory stress induce H3K9me3 loss at evolutionarily young LINE-1 and LTR elements in HSCs, resulting in persistent TE activation and long-term HSC dysfunction^13–16, 19, 20^. Here, we show that physiological aging induces remarkably similar alterations and that transient reprogramming largely prevents their establishment. Importantly, pharmacological inhibition of reverse transcriptase activity using lamivudine reduced DNA damage accumulation and improved HSC function in aged mice, indicating that TE activation contributes functionally to HSC decline rather than merely reflecting epigenetic disorganization.

Recent work has proposed that TE activation contributes to aging through the generation of dsRNAs and/or cDNA, activation of innate immune pathways, and chronic inflammatory signaling^14, 61, 62^. Consistent with these observations, we detected increased dsRNA accumulation and enhanced inflammatory signatures in aged HSCs, both of which were prevented by transient reprogramming. Our findings further suggest that an additional and potentially major consequence of TE activation in aged HSCs is the induction of persistent DNA damage. The beneficial effects of lamivudine on γH2AX accumulation strongly support a causal role for TE activity in age-associated genomic instability. Together with our previous work demonstrating the importance of young active L1Md elements in irradiation and chronic LPS-induced HSC DNA damage, and loss of HSC function^15, 19, 20^, these findings support a model in which TE activation may contribute to HSC aging through at least two non-exclusive mechanisms: activation of innate immune pathways through dsRNA sensing and induction of DNA damage through LINE-1 enzymatic activities.

AP-1 TFs also appear to play an important role in TE regulation. TEs that lose H3K9me3 or gain chromatin accessibility in aged HSCs were highly enriched for AP-1 motifs. Moreover, transient OSKM expression induced an early reduction in accessibility at AP-1-bound TE-associated regions in young mice, an effect that persisted with age. Importantly, pharmacological inhibition of AP-1 attenuated LPS-induced accumulation of dsRNA and γH2AX foci, as well as LINE-1 ORF1 protein expression, all of which were previously shown to result from L1Md derepression^20^. In contrast, AP-1 inhibition had no detectable effect in transiently reprogrammed HSCs, consistent with the notion that OSKM-mediated reprogramming had already prevented AP-1 activation and its downstream consequences on TE expression. These findings are reminiscent of recent observations showing that increased AP-1/c-Jun activity promotes chromatin opening at TE-containing genomic regions and TE mobilization in aged brain ultimately leading to impaired neurogenesis in Alzheimer’s disease^62, 63^. Together, these results suggest that AP-1-dependent regulation of TEs may represent a conserved mechanism linking stress exposure and aging to functional decline across distinct cell types.

Taken together, our findings support a model in which transient OSKM reprogramming resets the susceptibility of HSCs to age-associated stress rather than simply reversing isolated aging phenotypes. In this model, repeated inflammatory insults progressively establish AP-1-dependent chromatin states that promote TE derepression, DNA damage accumulation, and durable alterations in HSC function (Fig. 8K). More broadly, our data identify AP-1-associated chromatin remodeling as a candidate mechanism linking environmental stress exposure, TE activation, and stem cell aging. Given the emerging role of AP-1-positive inflammatory memory in HSCs in aging and clonal hematopoiesis, interventions aimed at preventing the establishment, maintenance, or reactivation of AP-1-dependent epigenetic programs may represent a promising therapeutic strategy to preserve HSC function and limit age-associated disease.

## Materials and methods

### Mouse strain, treatments and reconstitution experiments

C57BL/6J Rosa26^rtTA/+^; Col1a1^4F2A/+^ transgenic mice expressing a doxycycline (Dox)-inducible polycistronic transgene encoding the four Yamanaka factors (Oct4 [Pou5f1], Sox2, Klf4, and c-Myc [Myc] ; OSKM), separated by 2A peptide sequences and inserted into the *Col1a1* locus (hereafter referred to as 4TF mice) have been described previously^23^. OSKM expression was induced by treating 2–3-month-old mice with Dox (0.5 mg/mL) in drinking water for 16 days. Non-reprogrammed control mice (NT) carried the same genotype but did not receive Dox treatment. Owing to the ubiquitous and constitutive expression of rtTA, administration of Dox triggers widespread transcription of the OSKM polycistronic transgene. Mice were analyzed, 1 month after the end of the Dox treatment or at 20–24 months. To evaluate its ability to reverse aging-associated phenotypes, a separate cohort received Dox treatment at an advanced age (18–20 months) and was analyzed 4 months later. C57BL/6 CD45.2 or CD45.1 mice (7-8 weeks old) were from Envigo or Charles River laboratories, respectively. All the mice were housed in a specific pathogen-free environment. All procedures were reviewed and approved by the Animal Care Committee N°26 (APAFIS #35147-2022020315419972). For chronic inflammatory stimulation, 1 month after Dox treatment, 4TF mice received intraperitoneal injections of 6 µg LPS-B5 Ultrapure (Invivogen, tlrl-pb5lps) or an equivalent volume of PBS, 3 times a week for 4 weeks, as described^20^. Transplantation assays and HSC sorting were conducted 1 month later. For pharmacological inhibition of reverse transcriptase activity, aged wild-type (WT) C57BL/6 mice (18–20 months) received lamivudine (BOC Sciences) diluted in drinking water at a concentration of 2 mg/mL for 4 months. For reconstitution experiments, recipient mice were irradiated with 9.5 Gy with an X-ray irradiator (RX irradiator X-RAD 320). Engraftments were performed by retro-orbital injections. Depending on the experiment, recipient mice in competitive transplantation assays received a total of 1.5×10^6^ to 2×10^6^ BMMCs. For secondary transplantation experiment and serial BM transplantation experiment, recipient mice were transplanted with 2×10^6^ BMMCs. Peripheral blood was collected monthly after transplantation to assess donor chimerism by flow cytometry. Donor and recipient cells were distinguished based on the expression of CD45 congenic markers (CD45.1, CD45.2, or CD45.1/2, depending on the experiment).

### Cell harvest and culture

BM was harvested from femur, tibia and hip bones and stained with antibodies (Supplementary Table 13) allowing detection of lineage-negative (Lin^-^) Sca^+^ kit^+^ cells (referred to as LSK cells or HSPCs), HSCs (LSK-CD34^-^Flk2^-^), LT-HSCs (LSK-CD34^-^Flk2^-^CD150^+^CD48^-^), ST-HSC (LSK-CD34^+^Flk2^-^CD150^+^CD48^-^), LT-HSC MK (LSK-CD34^-^Flk2^-^CD150^+^CD48^-^CD41^+^); MPPs (LSK-CD34^+^Flk2^-^), and LMPPs (LSK-CD34^+^Flk2^+^). Samples were acquired using FACSCanto™ and Symphony™ A1 flow cytometers (BD Biosciences). Data were analyzed using FlowJo software (Tree Star Inc.), as described^15, 19, 20^. For HSC isolation, total BM was depleted of differentiated hematopoietic cells (lineage-positive; Lin+ cells) using Mouse Hematopoietic Progenitor Stem Cell Enrichment Set (BD Biosciences) and HSCs were sorted using FACSAria™ III, FACSAria Fusion (BD Biosciences), or Bigfoot (Thermo Fisher Scientific) cell sorters. For single cell liquid cultures, individual HSCs were sorted into round-bottom 96-well plates containing 50 µL of StemSpan™ SFEM medium (STEMCELL Technologies) supplemented with 1% penicillin/streptomycin (P/S), containing the following cytokines (all from Miltenyi Biotech): FTL3-Ligand (FTL3-L, 25ng/ml), stem cell factor (SCF, 50ng/mL) and thrombopoietin (THPO, 25ng/mL). Cells were cultured at 37°C with 5% CO2, and the number of cells per well was recorded after 2, 3, and 6 days. For each well, a transition from one cell to zero cells was scored as cell death, maintenance of a single cell was classified as quiescence, and the presence of more than one cell was considered indicative of proliferation. For dsRNA, LINE-1 ORF1p and γH2AX quantification in the presence of LPS, LSK cells were sorted into 200µL StemSpan supplemented with 1% P/S and FTL3-L (25ng/ml), SCF (50ng/mL) and interleukin-3 (IL3, 10ng/mL). After 24 hours of culture, LPS (10µg/mL) and / or AP-1 inhibitor T-5224 (10µM, HY-12270 MedChemExpress) were added and cells were incubated for an additional 48 hours at 37°C. HSCs (LSK, CD34^-^ Flk2^-^) were then sorted fixed and processed for IF.

### Clonogenic assays

250 HSCs were seeded in 500 µl methylcellulose medium (MethoCult™ GF M3434, STEMCELL Technologies) supplemented with 1% P/S in 24-well plates. Colonies were counted after 7 days at 37°C. Cells were then harvested, counted and replated in fresh methylcellulose medium at a density of 5,000 cells per well. In some experiments, LPS (10 µg/mL) and AP-1 inhibitor T-5254 (10 µg/mL) were added directly to the methylcellulose medium at the start of the culture. For lamivudine treatment, the cells were first cultured for 48 hours in liquid culture medium with cytokines, as above, in the presence or absence of 10 µM lamivudine (Sigma-Aldrich, 3TC-L1295) before being plated in methylcellulose.

### Immunofluorescence

IF analysis was performed as described^15^. Briefly, 2500-5000 HSCs were fixed with 4% paraformaldehyde (PFA) for 10 min at RT. After washing with PBS, the cells were resuspended into 200µL of PBS and cytocentrifuged at 500g for 5 minutes (Cellspin I Cytocentrifuge, Tharmac) on Poly-L-lysine coated slides. For yH2AX, H3K9me3, c-Jun, c-Fos and ORF1p staining, the cells were permeabilized for 5 min at RT with PBS containing 0.1% Triton, 0.1% Sodium citrate followed by 1 hour with a blocking buffer (PBS, 2% BSA, 2% FBS, 0.1% Triton). The slides were then incubated overnight at 4°C with anti-γHAX (1/2000, clone JBW301, EMD Millipore), anti-H3K9me3 (1/1000, Diagenode C1541005), anti-c-Jun (1/300, Cell Signaling 9165T), anti-c-Fos (1/500, Cell Signaling 2250T) and anti-ORF1p (1/750, Abcam ab216324) antibodies. For dsRNA staining, HSCs were permeabilized for 15 minutes with Triton X100 (0.5% in PBS) and blocking was done during 1 hour with 0.25% BSA at RT before staining with the anti-dsRNA antibody (1/150, clone J2, MABE1134, Sigma). Detection was performed using Alexa Fluor 555 or 488-coupled anti-mouse secondary antibody (1/600) (Invitrogen, A-21425). Slides were mount in a DAPI-containing mounting medium (DAPI Fluoromount-G Southern Biotech) and visualized using an SPE or Thunder confocal microscopes (Leica). Pictures were analyzed using ImageJ or CellProfiler softwares.

### CUT&Tag

H3K9me3 CUT&Tag was performed on 3,000 freshly sorted HSCs using the CUT&Tag-IT kit (Active Motif, #53160) according to the manufacturer’s instructions. Cells were incubated O/N with 0.5µg of H3K9me3 (C15410193-Diagenode). Transposed DNA fragments were amplified 18-fold by PCR using adapters supplied with the kit. PCR purification was carried out using the same Kit to remove remaining primers and large fragments. The quantity and quality of the libraries were assessed on Agilent 2100 Bioanalyzer (Agilent Technologies 50567-4626). Sequencing of the libraries was performed on the NovaSeq-6000 at Gustave Roussy (Illumina; 2 x50 bp).

### CUT&Tag analysis

Data integrity was first verified using md5sum (v8.22). Read quality was assessed with FastQC (v0.12.1) and MultiQC (v1.13), followed by adapter and quality trimming using Cutadapt (v0.6.10). Paired-end reads were aligned to the mm10 mouse reference genome using Bowtie2 (v2.4.1) with the parameters: --end-to-end --very-sensitive --no-mixed --no-discordant --phred33 -I 10 -X 700. Duplicate reads were removed using Picard (v2.26.9) with the parameters REMOVE_DUPLICATES=true and VALIDATION_STRINGENCY=LENIENT. Reads mapping to regular chromosomes were retained, and alignment quality scores were reset to 0 using SAMtools (v1.13). Aligned reads were processed using two complementary approaches. First, BAM files were sorted and indexed with SAMtools (v1.13), and peaks were called using SEACR (v1.3) in relaxed mode with a threshold of 0.01. Second, BAM files were downsampled to the minimum read count across samples, then sorted and indexed using SAMtools. These downsampled BAM files were used to quantify read counts within reproducible peaks across biological replicates. Downstream analyses were performed using custom R scripts (v4.1.2). A count matrix for union peaks was generated using the chromVAR package (v1.16.0). Differentially enriched peaks were identified using DESeq2 (v1.34.0), with significance thresholds of p-value < 0.05 and |log2FC| > 0. TE annotation was performed using the Bailly-Bechet database, as described^19^. Principal component analysis (PCA) was performed using normalized CUT&Tag counts from the 500 most variable regions. Hypergeometric tests were used to assess enrichment of H3K9me3within TE classes (LINE, LTR, SINE, and DNA transposons) relative to their genomic frequencies.

### Bulk RNA-seq

HSCs from individual mice were sorted directly into RLT buffer and stored at −80°C. Total RNA was extracted using the RNeasy Plus Micro Kit (Qiagen, #74034) according to the manufacturer’s instructions. To minimize genomic DNA contamination and improve TE quantification, an additional DNase treatment was performed using the gDNA Removal Kit (Enzo, ENZ-KIT136). RNA-seq libraries were generated using the SMARTer Stranded Total RNA-Seq Kit v3 – Pico Input Mammalian (Takara Bio). Library quality was assessed using an Agilent Bioanalyzer. Libraries from two independent experiments, comprising Y-NT, O-NT, and O-Dox samples, were pooled and sequenced together in a single run on a NovaSeq 6000 sequencer at the Gustave Roussy Genomics Core Facility using paired-end 100-bp sequencing. Libraries from a third independent experiment, comprising Y-NT and Y-Dox samples, were sequenced in a separate paired-end 100-bp run by BGI Genomics on a DNBSEQ-G400 instrument.

### Bulk RNA-seq analysis

Raw FASTQ files were assessed with FastQC (v0.12.1). Unique molecular identifiers (UMIs) were extracted using UMI-tools (v1.1.5), and adapter sequences were removed using Trim Galore (v0.6.10). Ribosomal RNA reads were filtered using SortMeRNA (v4.3.7). Reads were aligned to the GRCm38 mouse reference genome using STAR (v2.7.11b), and transcript-level quantification was performed with Salmon (v1.10.3). Transcript abundances were summarized to gene-level counts using tximport (v1.36.1). Differential gene expression analysis was performed using DESeq2 (v1.48.0), with differentially expressed genes (DEGs) defined as those with FDR < 0.1 and |log2FC| > 0.4. Gene Set Enrichment Analysis (GSEA) was conducted using clusterProfiler (v4.16.0) ^64^ on genes ranked according to −log10(p-value) × sign(log2FC), using the Hallmark and Reactome gene set collections. DEGs identified in the Old NT versus Young NT comparison were visualized using a heatmap generated with ComplexHeatmap (v2.24.0), applying k-means clustering (k = 5) to normalized, batch-corrected counts obtained from DESeq2. Over-representation analysis was subsequently performed for each cluster using clusterProfiler. Gene Set Variation Analysis (GSVA) ^65^ was performed using normalized, batch-corrected counts and the GSVA package (v2.2.0), together with published gene sets^33, 34, 53^, to estimate pathway activity across the different conditions.

TE expression was quantified using the SQuIRE pipeline^66^. RNA-seq reads were aligned to the mm10 reference genome using SQuIRE::Map, and expression levels at individual TE loci were quantified with SQuIRE::Count. TE copies with TPM ≥ 1, as well as L1Md elements longer than 4 kb displaying TPM ≥ 1 (average across biological replicates within each condition), were considered robustly expressed. Statistical differences between conditions were assessed using two-sided t-tests performed on replicate-level TPM values for individual TE copies.

### Single-cell RNAseq

scRNA-seq libraries were generated using the 10x Genomics GEM-X Universal 3’ Gene Expression v4 4-plex On-chip Multiplexing (OCM) platform (10x Genomics, Pleasanton, CA, USA) according to the manufacturer’s instructions. LSK Flk2^-^ HSPCs were sorted from 6 Y-NT, 6 Y-Dox, 4 O-NT and 4 O-Dox mice and pooled. Samples from Y-NT + Y-Dox and O-NT + O-Dox were multiplexed on 2 different GEM-X OCM chips using the on-chip multiplexing workflow. Using reverse transcription master mix, microfluidic partitioning was performed on a Chromium X instrument to generate Gel Beads-in-Emulsion (GEMs), enabling encapsulation of individual cells with uniquely barcoded gel beads. Within each GEM, polyadenylated mRNA molecules were captured by barcoded oligonucleotides containing a cell-specific barcode, a unique molecular identifier (UMI), and an oligo(dT) sequence for transcript capture. Following GEM generation, reverse transcription was carried out to synthesize barcoded first-strand cDNA. GEMs were subsequently broken, cDNA was purified and gene expression libraries were prepared from amplified cDNA through enzymatic fragmentation, end repair, A-tailing, adaptor ligation, sample indexing with dual-index primers, and final library amplification. After purification, final library quality and fragment size distribution were evaluated using an Agilent Fragment Analyzer and final library was sequenced on an Illumina NextSeq 2000 using paired-end sequencing with the read configuration recommended by 10x Genomics (28 bp for Read 1, 10 bp i7 index, 10 bp i5 index, and 90 bp for Read 2). 4070, 4186, 9372, 5740 cells, for Y-NT, Y-Dox, O-NT and O-Dox passed the quality control and were sequenced.

### Sc-RNA-seq analysis

Raw sequencing data were processed using the Cell Ranger pipeline (10x Genomics Cell Ranger, v9) and aligned to the mm10 reference genome. The cellranger count function identified 15,281 cells from old mice and 10,876 cells from young mice. All downstream single-cell data analyses were performed in R (v.4.3.1) using the Seurat package (v5.3.1) ^67^. Cells were filtered based on quality control metrics: > 1000 unique molecular identifiers (nCount_RNA > 1000), > 1000 detected genes (nFeature_RNA > 1000) and < 5% mitochondrial gene content (percent.mt < 5%). Following quality filtering, 11516 cells from old mice and 4021 cells from young mice were retained for further analysis. Standard normalization and scaling were applied while regressing out mitochondrial read proportions (percent.mt) and the difference between S phase and G2/M phase scores (CC.Difference = S.Score – G2M.Score). Experimental conditions were integrated using RunHarmony function of harmony package (v.1.2.3) ^68^. Unsupervised clustering was conducted on the top 20 principal components using the Smart Local Moving (SLM) algorithm at a resolution of 0.5, identifying 8 distinct cell clusters.

To characterize the molecular identity of each cluster, differential gene expression analysis was performed using the FindAllMarkers function (Wilcoxon rank-sum test; adjusted p < 0.05, log2Foldchange > 0.25). Cluster-specific marker genes were evaluated to annotate distinct hematopoietic stem and progenitor cell (HSPC) populations. Cluster annotations were further verified and confirmed by reference-based automated annotation using the SingleR package (v2.4.1), with standard murine reference datasets ^4, 8, 33, 69, 70^.

Differences in phenotypic cell populations (e.g., CD150 expression distribution across conditions) were evaluated using Fisher’s exact tests / Chi-square tests of independence on contingency tables. Pairwise post-hoc comparisons were conducted using the pairwiseNominalIndependence function with Benjamini-Hochberg False Discovery Rate (FDR) adjustment for multiple testing.

For each individual cell, the signature score was defined as the log-transformed mean normalized expression to of all genes in the given signature set. Score = log(mean(NormalizedCounts) +1). Single-cell signature activity was calculated using Seurat’s AddModuleScore function, which compute the average expression levels of each signature set on single cell level, subtracted by the aggregated expression of randomly sampled control feature sets. For signature score metrics (Z-scores, Module Score, and Log-Mean), pairwise comparisons were performed using two-sided Wilcoxon rank-sum tests (wilcox.test) with FDR correction. Adjusted p-values are reported as: ns (p > 0.05), * (p < 0.05), **(p < 0.01), ***(p < 0.001), and ****p < 0.0001).

Pathway enrichment analysis was performed by clusterProfiler (v.4.10.0). The MSigDB Hallmark(H) database were utilized, pathways with an adjusted FDR < 0.05 were considered statistically significant.

### ATAC-seq

ATAC-seq was performed on 3,000 freshly sorted HSCs using the Active Motif ATAC-Seq Kit (#53150) according to the manufacturer’s recommendations. Library quality was assessed using an Agilent Bioanalyzer, and paired-end sequencing (2 x 50 bp) was performed on an Illumina NovaSeq 6000 platform.

### ATAC-seq analysis

Paired-end reads were trimmed using Trim Galore (v0.6.10) and aligned to the mm10 genome using Bowtie2 (v2.3.5.1) with the parameters --end-to-end --very-sensitive --no-mixed --no-discordant -- phred33 -X 2000. Duplicate reads were marked and removed using Picard (v2.23.8). SAMtools (v1.10) was used for filtering, downsampling, and indexing aligned reads. Blacklisted regions and reads mapping outside standard chromosomes were excluded. Peaks were called using MACS2 (v2.2.7.1) with the parameters --nomodel --shift -100 --extsize 200 --broad. Differentially accessible regions (DARs) were identified using DiffBind (v3.8.4). Regions with FDR < 0.1 were considered significantly differentially accessible. Heatmaps were generated from z-score-scaled normalized read counts using ComplexHeatmap (v2.22.0). DARs were annotated using HOMER’s annotatePeaks.pl script with the - size given option to preserve original peak coordinates. Each DAR was assigned to its nearest gene, and pathway over-representation analyses were performed to identify enriched biological processes. DARs identified in the Old NT versus Young NT comparison were clustered using k-means clustering (k = 7) to identify regions displaying reduced accessibility in Young Dox relative to Young NT samples. Over-representation analyses were subsequently performed on the nearest genes associated with these clusters.

Motif enrichment analyses were conducted using HOMER’s findMotifsGenome.pl script (v4.11) with the -size given parameter. Only motifs with enrichment p-values < 1 × 10⁻³⁰ were retained. BAM files were converted to bigWig format using bamCoverage from DeepTools (v3.5.5), and heatmaps were generated using plotHeatmap. For footprinting analysis, for each condition, a peak set was generated by merging peaks and retaining only regions consistently detected across all biological replicates. Consensus peak sets were then generated by merging condition-specific peak sets. Footprinting and differential TF binding analyses were performed using TOBIAS^39^ (v0.14.0), using consensus peak sets and condition-merged BAM files as input. AP-1-bound DARs were identified from TOBIAS outputs by selecting regions containing footprints associated with both FOS and JUN family motifs. TOBIAS was additionally run across all conditions simultaneously using a global consensus peak set, allowing direct comparison of TF binding activity across all samples.To quantify TE accessibility, DiffBind analyses were performed using the TE annotation database rather than MACS2 peak calls, as described^19, 20^. TOBIAS footprinting analyses were restricted to TE-overlapping accessible regions, and AP-1-bound TE-associated regions were identified based on footprints matching FOS/JUN family motifs.

### Quantitative RT-PCR

RNAs were reverse transcribed using the HiScript III RT SuperMix for qPCR (+gDNA wiper) kit (Vazyme R323-01). 1.25 µL of cDNA was pre-amplified for 15-18 cycles in a multiplex reaction using Preamp Master-Mix solution (ref. 100-5580, Fluidigm) and primer mix (200 µM of each primer; Supplementary Table 14), as described ^15^. Quantitative PCR (qPCR) was performed using LUNA Universal qPCR Master Mix (New England Biolabs) on the 7500 real-time PCR machine (Applied Biosystems). Data were normalized with the mean expression of HPRT.

### Statistics

Statistical analyses were performed using GraphPad Prism software (version 8.0). The results are displayed as the means and SEM. The value of *, P < 0.05 was considered as significant, and **, P < 0.01 or ***, P < 0.001 as highly significant.

## Data availability

The dataset generated from the bulk RNAseq, single-cell RNA-seq, CUT&Tag and ATAC-seq experiments for Figures 3, 4, 5, 6, 7 are available in ArrayExpress accession E-MTAB-17474 and E-MTAB-17476; E-MTAB-17477; E-MTAB-17534; E-MTAB-17478, respectively.

## Acknowledgments

We thank animal facility from Gustave Roussy and UMS44 – IBVB, Hôpitaux Bicetre & P. Brousse, the Imaging and Cytometry and the Genomic Platforms for cell sorting, confocal analyses, and for for CUT&Tag, and ATAC and sc-RNA sequencing, respectively. This work was supported by INSERM and grants from Ligue Nationale Contre le Cancer (LNCC, Equipe labellisée EL2020), ARC Foundation (n° ARCPGA2023110007352_7985) and Agence Nationale de la Recherche (ANR-23-CE14-0017) to F.P. FACS analysis was supported by DIM-ITAC Région Ile de France to F.P. A.P is recipient of a fellowships from the Ministère de l’Enseignement Supérieur de la Recherche et de l’Innovation and from the Société Française d’Hématologie. A.P received fundings from the Ministère de l’Enseignement Supérieur de la Recherche et de l’Innovation and from the French Society of Hematology. M. B and M.Y are supported ANR-23-CE14-0017 to F.P. H.A.O is supported by ARC foundation (n° ARCPGA2023110007352_7985 to F. P.

## Author contributions

A.P. performed all experiments and analyzed the results. M.B. participated to blood and BM sampling, reconstitution experiments and FACS analyses. H-A.O., R.C., M.Y and E.Z. performed bioinformatic analyses; N.D. performed sc-RNA sequencing; T-H T and C.M.S analyzed single cell RNA-seq data; O. M. and J-M. L provided the 4TF strain; F.P. designed and supervised the study, analyzed the results and wrote the manuscript.

