## Supplementary figures for "Transient reprogramming limits AP-1-associated chromatin opening and transposable element activation to preserve hematopoietic stem cell function during aging and stress"

**Extended Fig. 1**

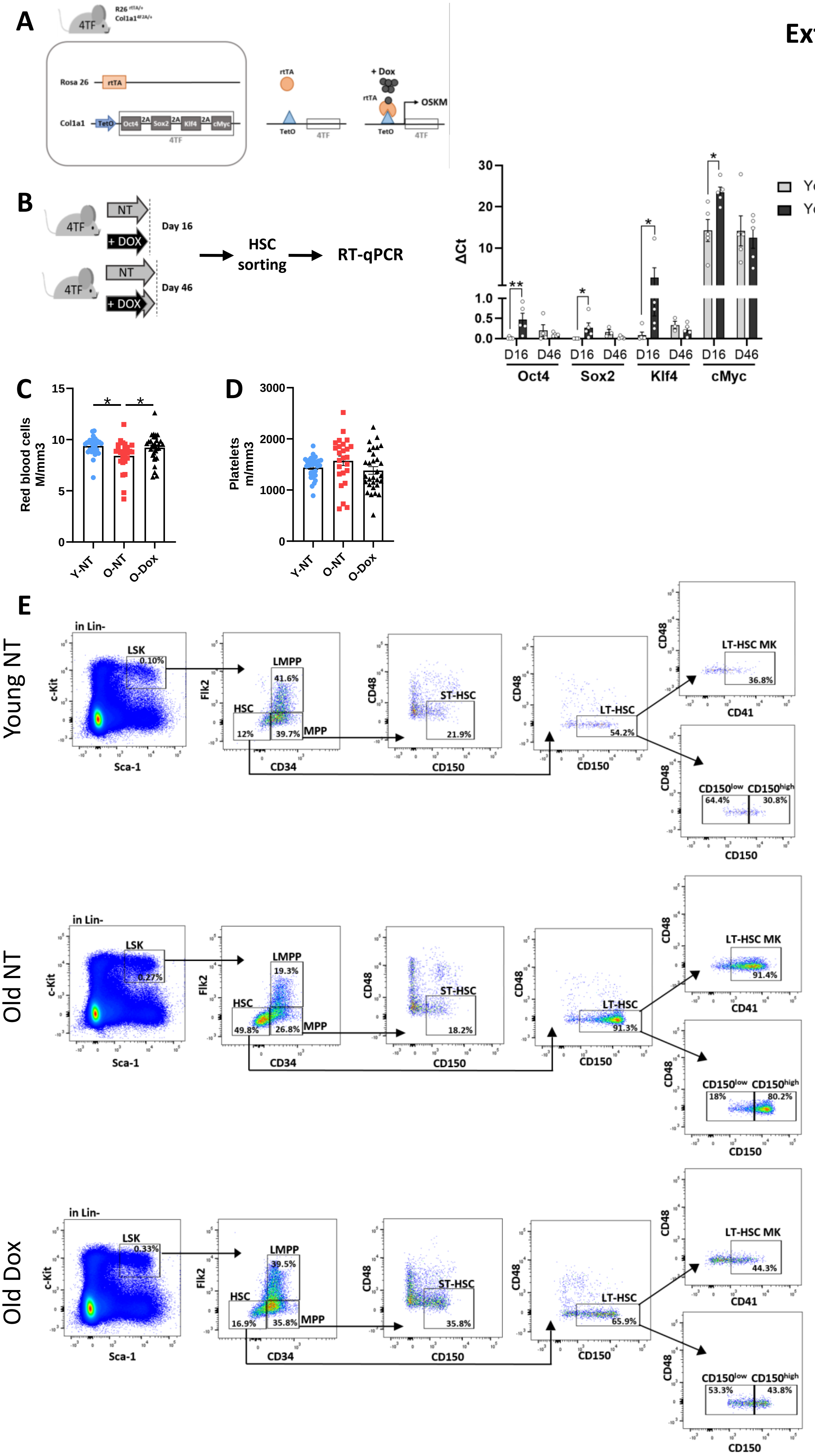

**Extended Figure 1 (related to Fig. 1)**

**(A)** Rosa26 $^{rtTA/+}$; Cola1$^{4TF2A/+}$ heterozygous 4TF mouse model expressing a polycistronic cassette containing OSKM sequences into the ubiquitous Collagen 1 locus and carrying the rtTA trans-activator into the Rosa26 locus. In the presence of Dox, rtTA is ubiquitously expressed and able to bind TetO promotor allowing OSKM expression.

**(B)** *Oct4*, *Sox2*, *Klf4* and *Myc* mRNA expression levels measured by RT-qPCR at the end of the Dox treatment (Day 16) and 1 month after Dox treatment arrest (Day 46), compared to NT mice. DCt values normalized to *Hprt* levels. Means +/- SEM. Unpaired t-test. * p-val <0.05.

**(C, D)** Red blood cell (C) and platelet counts (D) in the peripheral blood of Y-NT, O-NT and O-Dox mice. Each dot represents a mouse. Means +/- SEM. One-way ANOVA with multiple comparison test. Adjusted p-values: * p<0.05.

**(E)** Representative FACS gating strategy for the analysis of HSCs (LSK CD34$^-$ Flk2$^-$), LT-HSCs (LSK CD34$^-$ Flk2$^-$ CD150$^+$ CD48$^-$), myeloid-biased LT-HSCs (LSK CD34$^-$ Flk2$^-$ CD150$^{high}$ CD48$^-$), lymphoid-biased LT-HSCs (LSK CD34$^-$ Flk2$^-$ CD150$^{low}$ CD48$^-$), megakaryocytic-biased LT-HSCs (LSK CD34$^-$ Flk2$^-$ CD150$^+$ CD48$^-$ CD41$^+$), lymphoid multipotent progenitor (LMPP) (LSK CD34$^+$ Flk2$^+$), multipotent progenitors (MPP) (LSK CD34$^+$ Flk2$^-$) and ST-HSCs (LSK CD34$^+$ Flk2$^-$ CD150$^+$ CD48$^-$) for Y-NT, O-NT and O-Dox mice.

### Extended Fig. 2

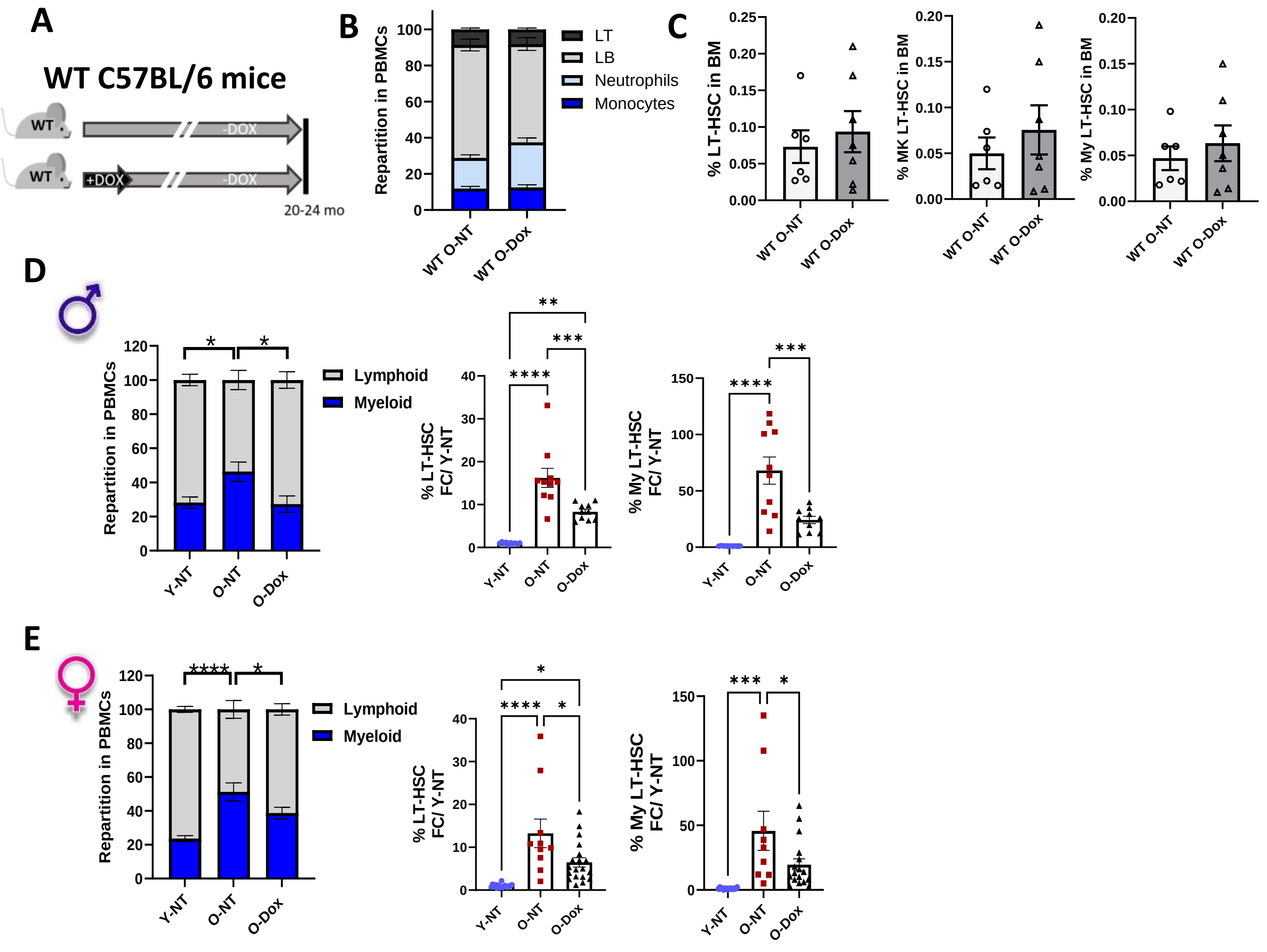


**Extended Figure 2 (related to Fig. 1)**

**(A-C)** Young (2-3 months) non-transgenic (WT) B6 mice were treated with Dox or left untreated. Blood (B) and BM LT-HSC (C) cell composition was analyzed by FACS at 20-24 months of age. NT (n=6), Dox (n=7) mice. Means +/- SEM. Two-way ANOVA with multiple comparison test (B) or unpaired t-test (C). **(D, E)** Blood and BM analysis in Y-NT, O-NT and O-Dox 4TF males (D) and females (E) separately. One dot represents a mouse. Data are normalized to Y-NT group. Means +/- SEM. One-way ANOVA with multiple comparison test, adjusted p-values are reported as: * p<0.05; ** p<0.01; *** p<0.001; **** p<0.0001

Extended Fig. 3

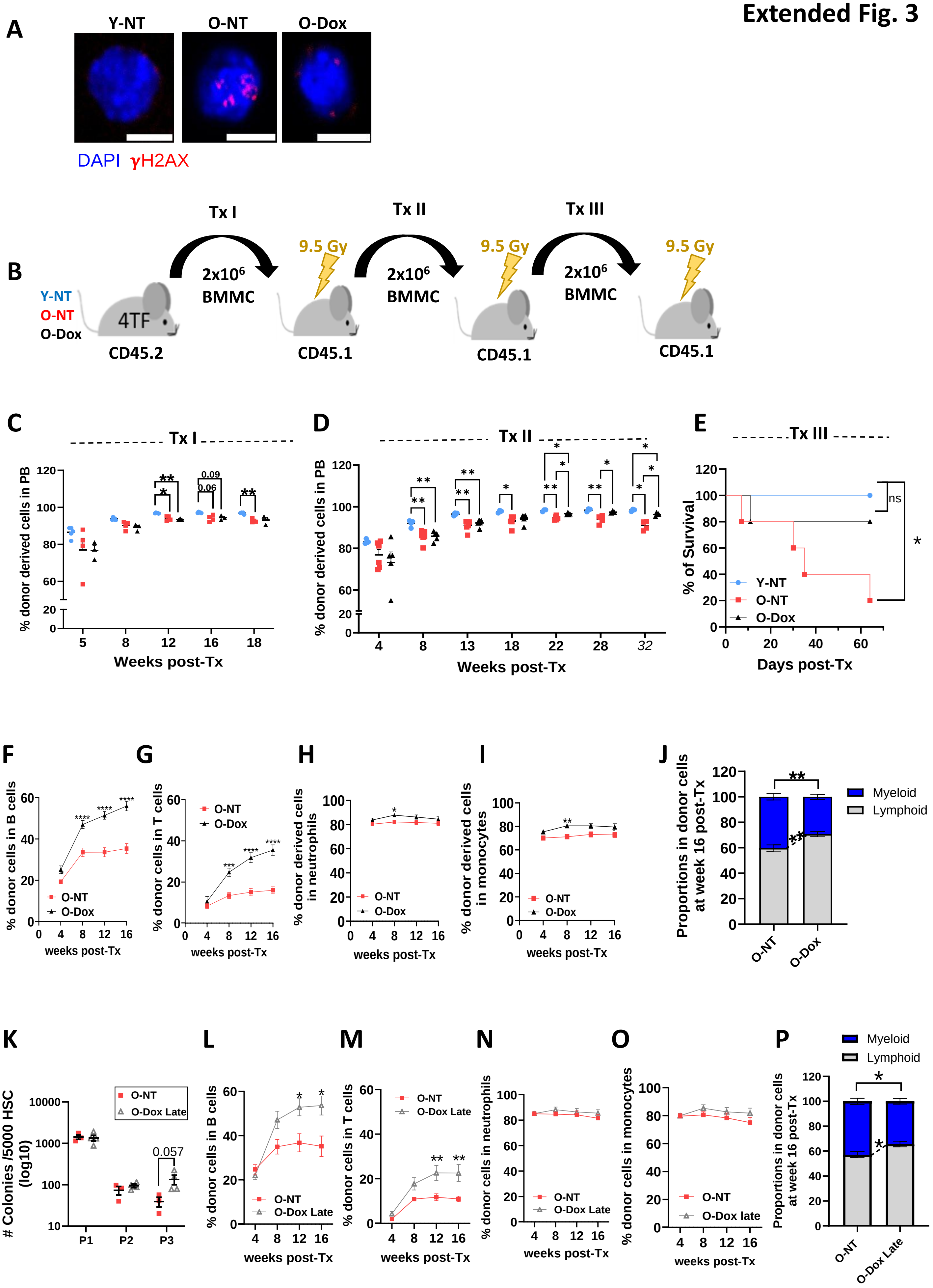

**Extended Figure 3 (related to Fig. 2)**

**(A)** Representative pictures of γH2AX IF presented in Fig. 2F. Scale bars: 5µm.

**(B-E)** Non-competitive serial transplantations of CD45.2$^{+}$ Y-NT, O-NT and O-Dox BM cells into young lethally irradiated CD45.1$^{+}$ recipients. (B) Experimental design. (C, D) Percentages of donor-derived cells in the blood of primary (C) and secondary recipients (D) over time. Each dot represents an individual mouse. Two-way ANOVA with multiple comparison test. Adjusted p-values are reported as: * $p<0.05$; ** $p<0.01$. (E) Survival of the third recipients during the first 60 days following transplantation. Log-rank Mantel Cox test.

**(F-I)** Percentage of donor-derived cells in B (F) and T (G) lymphocytes, neutrophils (H) and monocytes (I) of recipients in the BM competitive transplantation presented in Fig. 2G. Means +/ SEM.

**(J)** Reconstitution bias illustrated as the percentages of myeloid (monocytes and neutrophils) and lymphoid (B and T cells) cells within donor derived cells in the blood of recipient 4 months following competitive transplantation with O-NT or O-Dox BM cells. Two-way ANOVA, adjusted p-values are reported as: * $p < 0.01$.

**(K)** Number of colonies generated from 5000 HSCs from O-NT and O-Dox late mice after plating in methylcellulose at passages 1, 2 and 3. Each dot represents an individual mouse. Means +/ SEM. Multiple t tests. *$p<0.05$; **$p<0.01$.

**(L-O)** Percentages of donor derived cells in B (K) and T (L) lymphocytes, neutrophils (M) and monocytes (N) of recipients after competitive transplantation with O-NT or O-Dox late BM cells (as presented in Fig2 L).

**(P)** Reconstitution bias illustrated as the percentages of myeloid (monocytes and neutrophils) and lymphoid (B and T cells) cells within donor derived cells in the blood of recipient 4 months following competitive transplantation with O-NT or O-Dox late BM cells. Two-way ANOVA, adjusted p-values are reported as: * $p < 0.05$.

**Extended Fig. 4**

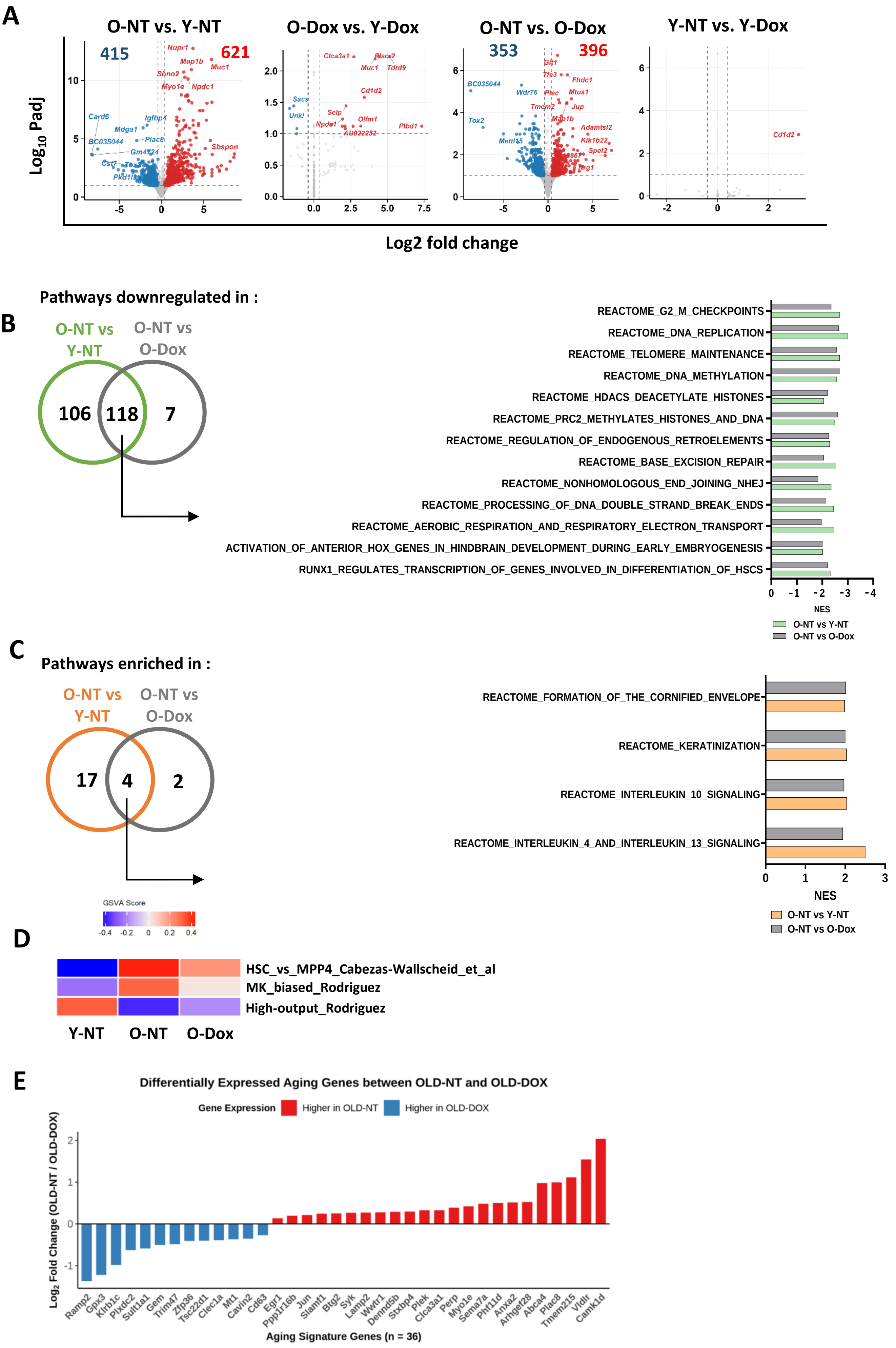

**Extended Figure 4 (related to Fig. 3)**

**(A)** Volcano plots showing up- (red) and down- (blue) regulated (FDR < 0.1; log2FC $\geq$ 0.4) and non-significant DEGs (grey) identified by bulk RNA-Seq for the indicated comparisons.

**(B, C)** Venn diagram and normalized enrichment scores (NES) of representative REACTOME gene sets (FDR< 0.05) commonly downregulated (B) or up-regulated (C) in O-NT samples compared to Y-NT and O-Dox. O-NT vs Y-NT: green and orange circles; O-NT vs O-Dox (grey circles).

**(D)** Gene set variation analysis (GSVA) measuring pathway activity using published RNA-Seq data sets in Y-NT, O-NT, O-Dox samples.

**(E)** Comparison of differentially expressed aging genes between O-NT and O-Dox LT-HSCs in single cell RNA-Seq. For each DEG, the Log2 Fold Change in gene expression between O-NT vs O-Dox was calculated using expression values from individual LT-HSCs.

**Extended Fig. 5**

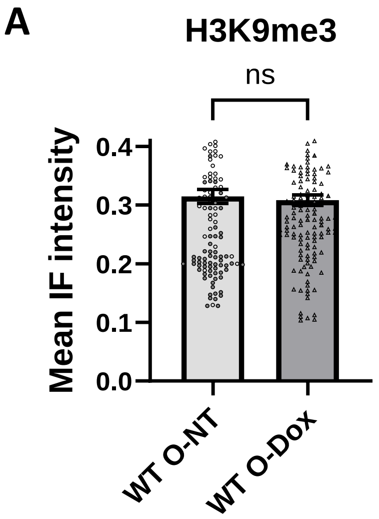


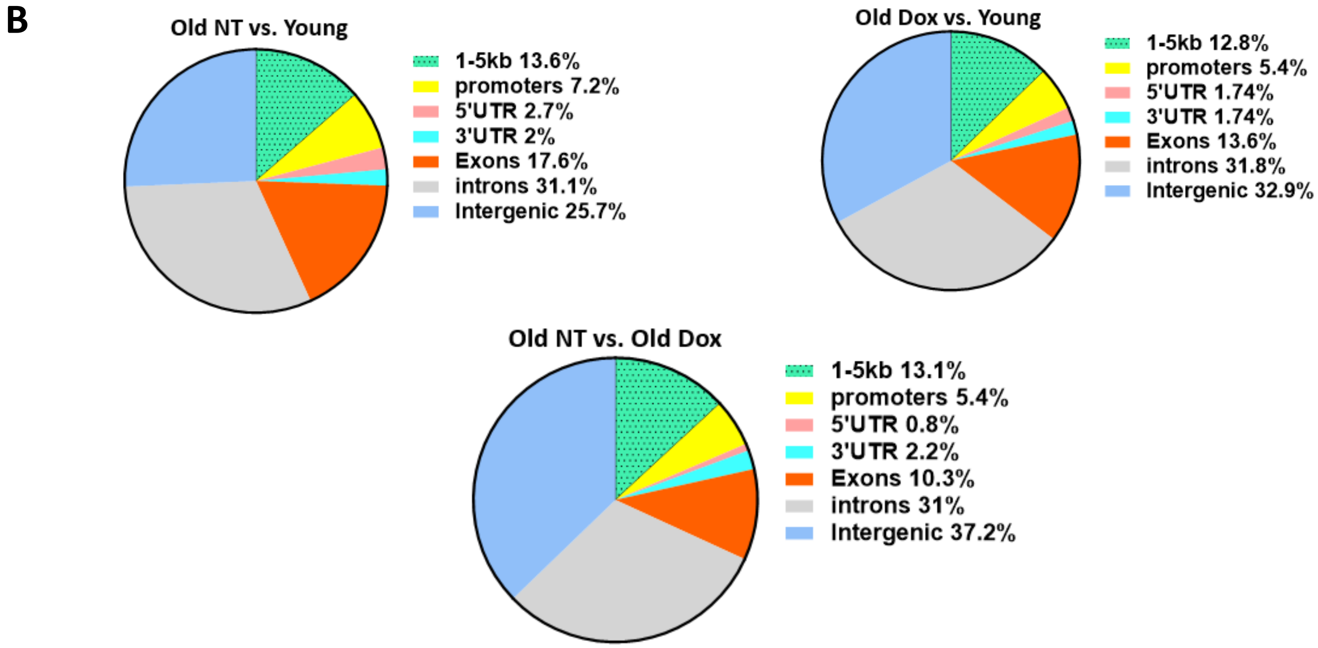


**Extended Figure 5 (related to Fig. 4)**

(**A**) IF analysis of H3K9me3 signal in HSCs from O-NT and O-Dox WT B6 mice. Results from at least 50 HSCs per mouse. 2 mouse per group.

**(B)** Distribution of H3K9me3 differential peaks genomic repartition for the indicated comparisons.

Extended Fig. 6

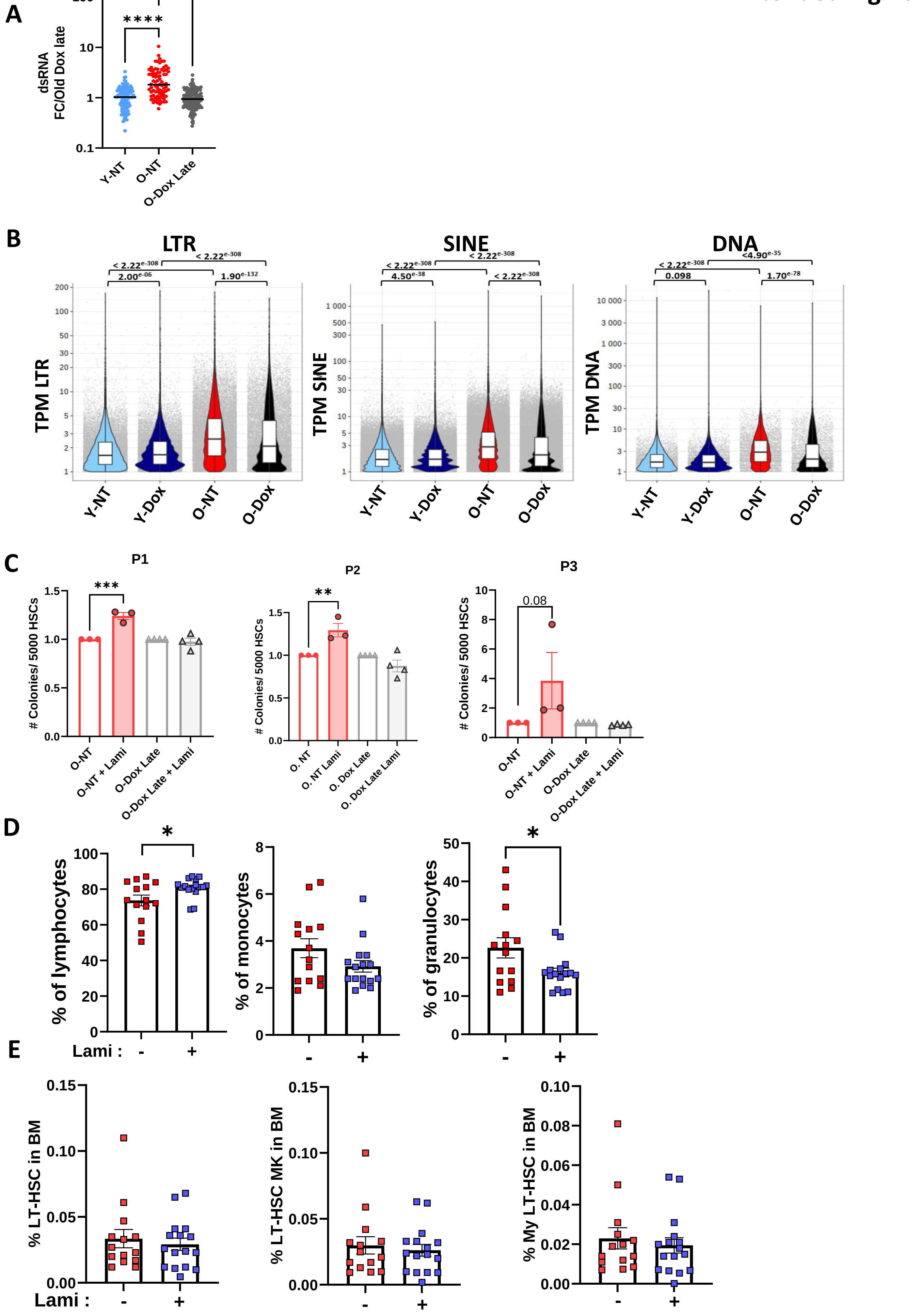

**Extended Figure 6 (related to Fig. 5)**

**(A)** Representative images and quantification of dsRNA IF in HSCs from Y-NT (n=2), O-NT (n=2) and O-Dox Late (n=5) mice. Results from at least 50 HSCs per mouse. Each dot represents one HSC. Means +/- SEM. One-way ANOVA. Adjusted p-values reported as **** p>0.0001. Scale bars represent 5µm.

**(B)** Violin plots showing the expression of LTR, DNA and SINE family TEs copies in bulk RNA-Seq data from HSCs from the indicated groups. One-way ANOVA, adjusted p-values are shown.

**(C)** Clonogenic activity of HSCs from O-NT and O-Dox late mice treated or not for 48h in vitro with lamivudine (lami, 10µM) at passages 1, 2 and 3. Paired t-test. ** p-val <0.01 and *** p-val <0.001.

**(D, E)** Blood cells counts (D) and FACS analysis of LT-HSCs (E) in the BM from aged WT mice treated or not with Lami *in vivo* for 4 months, as in Fig. 5E. Each dot represents an individual mouse. Means +/- SEM. Unpaired t- test. * p-val <0.05.

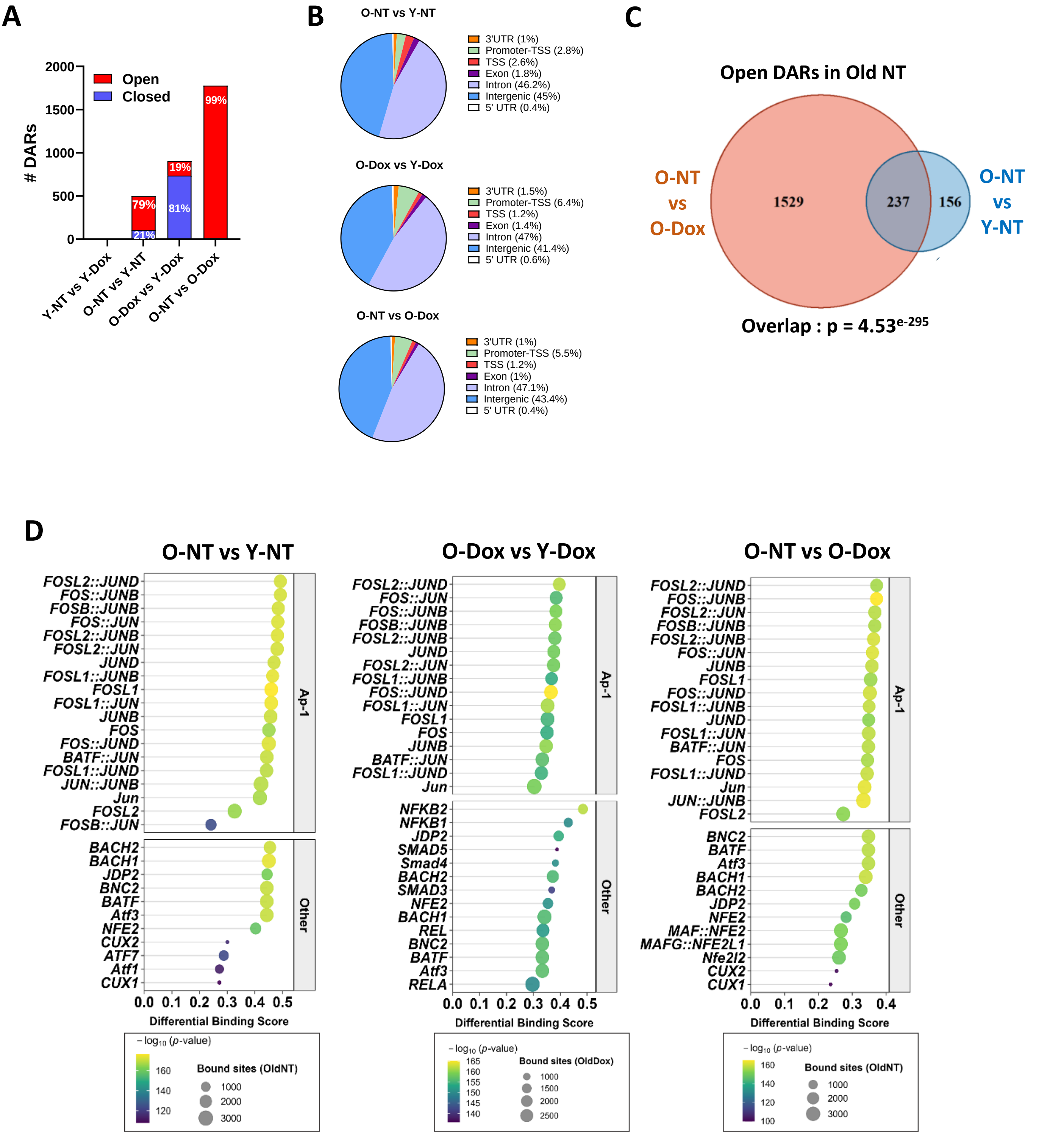


**Extended Figure 7 (related to Fig. 6)**

**(A)** Number of ATAC-seq open DARs (red) and closed DARs (blue) for the indicated comparisons.

**(B)** Genomic distribution of DARs for the indicated comparisons.

**(C)** Overlap of open DARs identified in the O-NT vs Y-NT and O-NT vs O-Dox comparisons. All peaks (significant and non-significant) are used as background and the statistical value of overlap is computed with a hypergeometric test.

**(D)** Top 30 HOMER TF binding site motifs identified in O-NT vs Y-NT (left), O-Dox vs Y-Dox (middle) and O-NT vs O-Dox (right) DARs. TFs are ranked by binding score. AP-1 TFs are grouped on the top panels.

### Extended Fig. 8

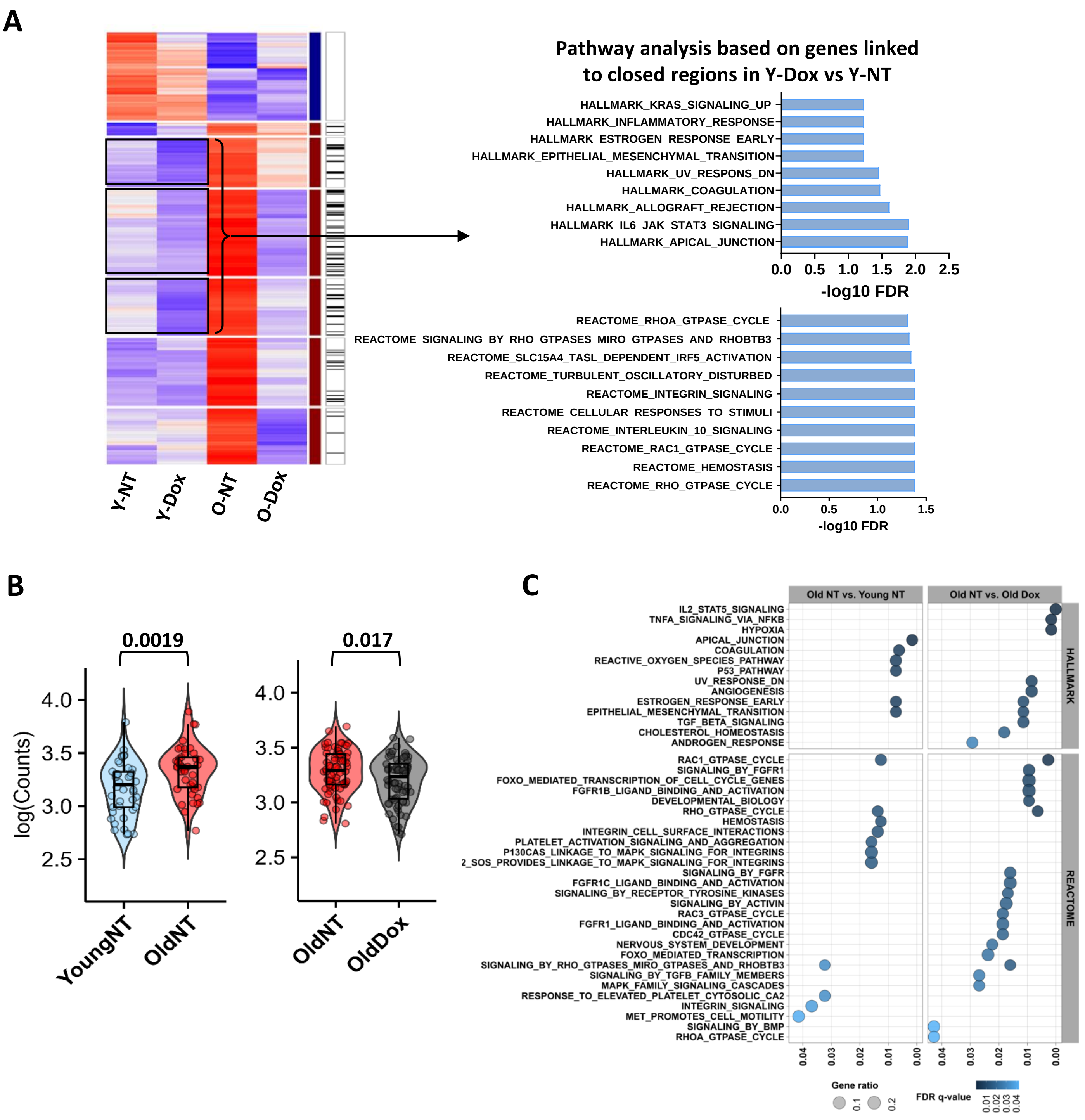


**Extended Figure 8 (related to Fig. 7)**

**(A)** GSEA pathway analysis performed with the list of genes associated to the ATAC-seq regions closed in Y-Dox vs Y-NT and open in Y-NT vs O-NT (black boxes).

**(B)** Violin plots showing the log-transformed RNA-seq read counts at DEGs associated with O-NT vs Y-NT (left) and O-NT vs O-Dox (right) ATAC-seq open DARs. One dot represents a single DEG. Unpaired t-test.

**(C)** Bubble plots presenting the pathways linked to DAR-closest DEGs in B for the indicated comparisons.

### Extended Fig. 9

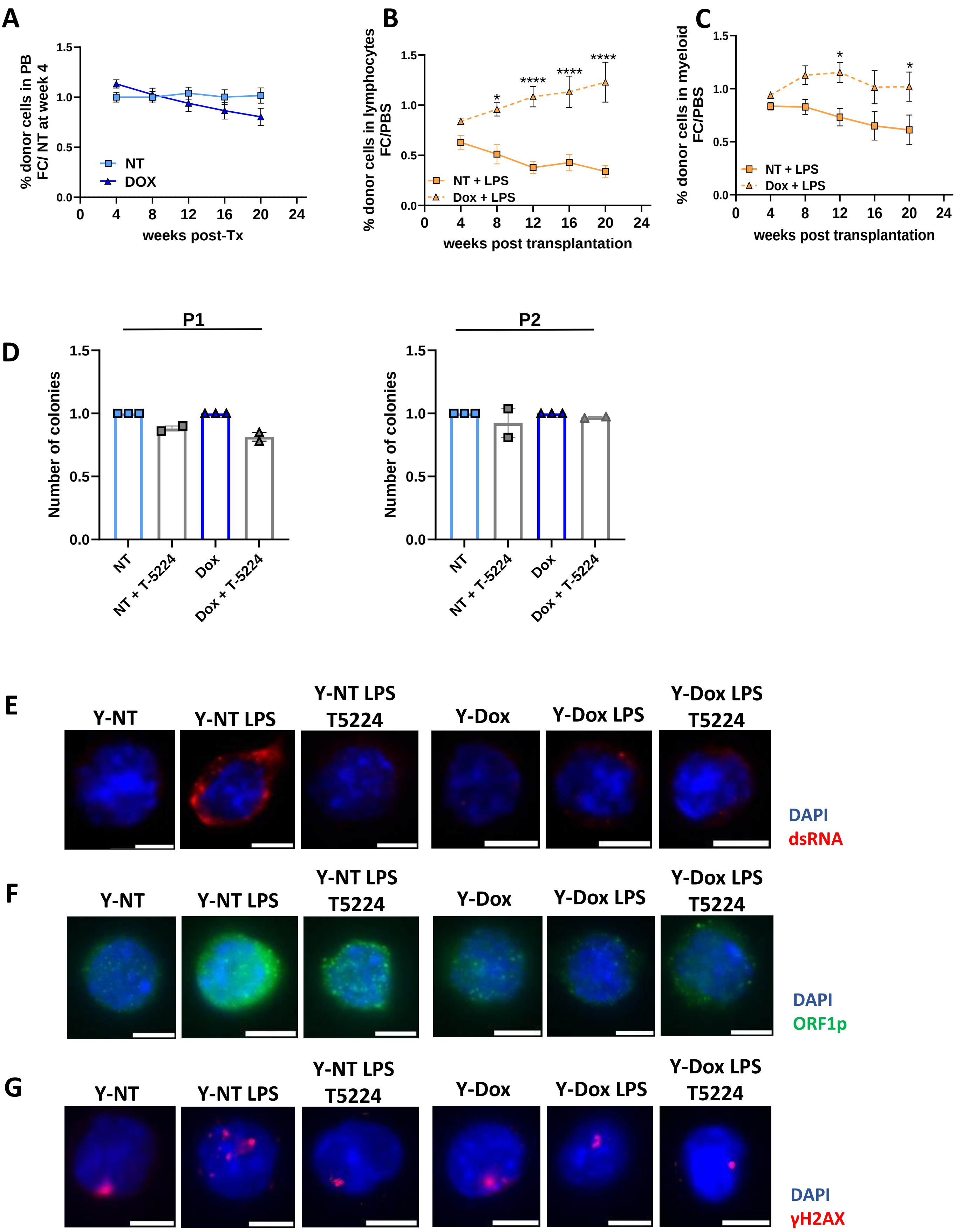

**Extended Figure 9 (related to Fig. 8)**

**(A)** Percentages of donor-derived cells in the peripheral blood of recipient mice engrafted with BM from Y-NT and Y-Dox PBS controls (Fig. 8A). Results presented as a fold change compared to NT group at week 4 post-Tx.

**(B, C)** Percentages of donor-derived cells in the lymphoid (B) and myeloid (C) compartments of mice engrafted with Y-NT or Y-Dox mice treated with PBS or LPS. Results are shown as the fold-change of LPS over PBS control group. Means +/- SEM, n=10 mice per group from 2 independent experiments. Two-way ANOVA with multiple comparisons. Adjusted p-values are reported as: * $p<0.05$, ** $p<0.01$, *** $p<0.001$.

**(D)** Number of colonies generated from 5000 HSCs from Y-NT and Y-Dox mice in the presence or absence of AP-inhibitor alone (10 µM). (E-G) Representative pictures of dsRNA (E) ORF1p (F) and γH2AX (G) IF presented in Fig. 8H, 8I and 8J, respectively. Scale bars: 5µm.
